# Integrative multi-omics analysis of *Polypterus* fin regeneration reveals shared and derived fin and limb regeneration mechanisms

**DOI:** 10.1101/2025.03.12.642910

**Authors:** Josane F. Sousa, Gabriela Lima, Louise Perez, Hannah Schof, Igor Schneider

## Abstract

Comparative studies of salamander limbs and fish fins may reveal shared components of an ancestral vertebrate appendage regeneration toolkit. However, traditional fish models, such as zebrafish, regenerate only dermal fin rays, which are neither homologous to limbs nor as cellularly complex. To address this, we employed a multi-omics approach leveraging the regenerative capacity of *Polypterus senegalus*, a fish species capable of full fin regeneration. Single-nucleus RNA-sequencing revealed extensive erythrocyte and myeloid cell infiltration during *Polypterus* fin regeneration. Spatial transcriptomics identified key cellular contributors to the blastema, while cross-species comparisons with axolotl limbs uncovered both common and divergent expression patterns of wound epidermis and blastema markers. Fin amputation triggered a conserved injury response, including sequential activation of pro- and anti-inflammatory programs and DNA damage repair in cycling cells. Hypoxia responses involved *hif1a* modulation, consistent with other regenerative models, but uniquely included the expression of *hif4a* in erythrocytes and the upregulation of a myoglobin gene in the wound epidermis. Furthermore, genome-wide chromatin profiling identified candidate regeneration-responsive elements. Together, these findings establish *Polypterus* as a powerful comparative model for identifying shared and derived mechanisms of limb and fin regeneration.

## Introduction

Among limbed vertebrates (tetrapods), salamanders are the only clade capable of regrowing severed limbs as adults. This regenerative capacity may have been inherited by tetrapods from their fish ancestors, retained in modern salamanders, partially preserved in frogs (limited to larval stages), but entirely lost in amniotes (Fig. 1a). Supporting this hypothesis, fossil evidence suggests that limb regeneration predates the origin of stem salamanders^1,2^. A phylogenetic survey of complete paired fin regeneration—defined as regeneration following amputation at the proximal fin endoskeleton— demonstrated that this ability is present across species representing all major bony fish clades^3,4^, including zebrafish larvae^5,6^. Furthermore, decades of comparative studies using the zebrafish tail fin as a model system have revealed commonalities between limb and fin ray regeneration programs^7^, even extending to cell-type-specific genetic and epigenetic programs^8^. Altogether, these findings support a shared evolutionary origin of limb and fin regeneration and underscore the importance of comparative studies as a powerful framework for identifying the core cellular and molecular components underlying vertebrate appendage regeneration.

**Fig.1.**
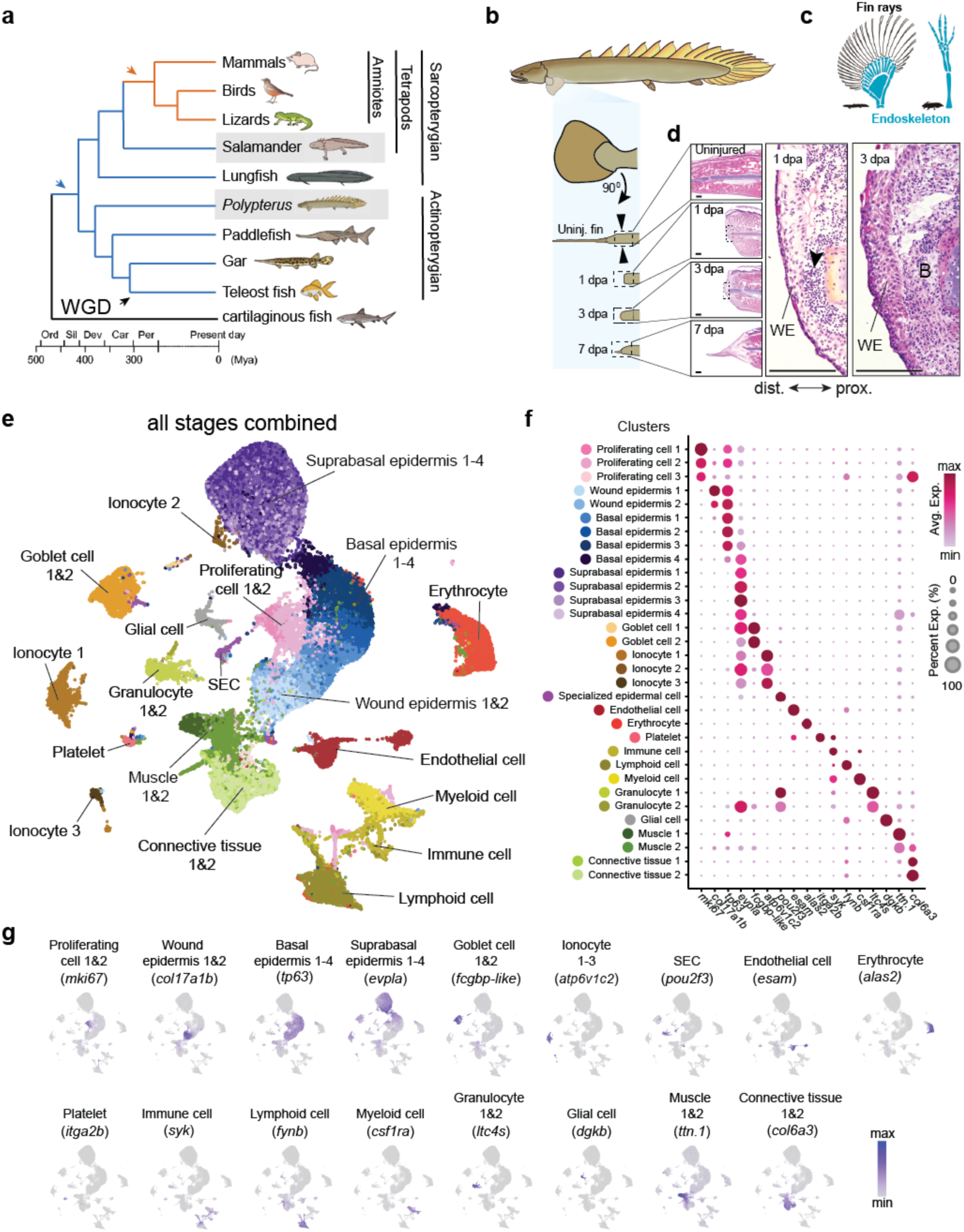
SnRNA-seq uncovers cellular diversity of *Polypterus* fin regeneration at homeostasis and regrowth. **a** Phylogeny of major vertebrate clades showing absence (black line), presence (blue lines), or absence (orange lines) of paired appendage regeneration; arrows indicate gain (blue) or loss (orange of regenerative capacity) and whole genome duplication event (black). **b** Proximal amputation of the pectoral fin of *Polypterus*; black arrowheads flanking uninjured fin drawing denote amputation site. **c** *Polypterus* fin rays (black) and endoskeletal elements (blue); axolotl limb endoskeletal elements (blue), and (**d**) histological sections of stages assayed in this study; WE, wound epidermis; black arrowhead in 1 dpa histological pane indicates immune cell infiltration; B, blastema. **e** Uniform manifold approximation and projection (UMAP) of nuclei from all stages (uninjured, 1 dpa, 3 dpa, and 7 dpa) identified 32 distinct clusters. **f** Heatmap of representative gene markers of major cell types identified in our dataset. **g** UMAP of representative gene markers of major cell types shown in (**e**). Scale bars = 100 μm.

While the adult zebrafish serves as a valuable model for comparative and functional studies in fin regeneration, its regenerative capacity is limited to the distal portion of the fin, which consists of dermal fin rays^5^ (Fig. 1c). These structures have no direct homologs in limbs^9^ and lack the cellular complexity and diversity observed during limb regeneration^8^. Additionally, the teleost-specific whole-genome duplication (WGD) event presents challenges for genetic comparisons between zebrafish and salamanders, complicating the identification of conserved regenerative mechanisms^10,11^. To address these limitations, we have established the Senegal bichir (*Polypterus senegalus*) as a model, leveraging its remarkable regenerative abilities. *Polypterus*, a non-teleost ray-finned fish (actinopterygian), diverged from teleosts prior to the teleost-specific WGD (Fig. 1a), maintaining a 1:1 gene orthology with tetrapods^12^. Unlike zebrafish, *Polypterus* can readily regenerate entire fins as adults, including the proximal endoskeleton, which shares deep homology with limb bones ^13,14^(Fig. 1c). Furthermore, our previous studies identified signaling pathways commonly upregulated in both salamanders and *Polypterus*^4^, highlighting its potential as a powerful model for evolutionarily informed comparative studies of vertebrate appendage regeneration.

To comprehensively identify the core cellular, genetic, and gene regulatory components of *Polypterus* fin regeneration, we employed single-nucleus RNA-sequencing (snRNA-seq). This approach allowed us to characterize the cell types present during homeostasis and across different stages of fin regeneration, enabling a comparative analysis of their similarities and differences with those previously described in axolotls. *Polypterus* fin regeneration was accompanied by increased cell proliferation and massive infiltration of erythrocytes and immune cells. Integration of snRNA-seq and spatial RNA-sequencing (spatial RNA-seq) revealed blastema heterogeneity, wound epidermis markers, the sequential activation of pro- and anti-inflammatory genetic programs, DNA damage response, and deployment of conserved and derived mechanisms of response to hypoxia. Finally, using the assay for transposase-accessible chromatin with high-throughput sequencing (ATAC-seq), we identified features of the epigenetic programs of fin regeneration.

## Results

### The cellular landscape of *Polypterus* fin regeneration

To assess fin regeneration at the single-cell level, we selected timepoints representative of homeostasis (the uninjured proximal segment of the pectoral fin) and regeneration at 1, 3, and 7 day-post amputation (dpa) (Fig. 1b). Fin amputation at the level of the endoskeletal elements resulted in complete wound closure and presumptive immune cell infiltration at 1 dpa (Fig. 1d). At 3 dpa, the wound epidermis was multilayered, containing distinct basal and suprabasal layers, and subjacent mesenchymal cells forming the presumptive blastema. At 7 dpa, the wound epidermis extended distally, forming a dorso-ventrally constricted outgrowth in which fin rays will form.

Using combinatorial barcoding technology (Parse Biosciences), we profiled over 63,000 nuclei from fins at homeostasis and during regeneration in biological replicates. Trailmaker (Parse Biosciences, 2024) was used to assess quality control metrics (Supplementary Fig. 1a) and complete our snRNA-seq data analysis. Unbiased clustering of all cells from all timepoints revealed 32 transcriptionally distinct populations (clusters) (Fig. 1e and Supplementary Fig. 1b). Cluster annotation based on transcriptional profiles identified multiple cell populations, including proliferating cells, muscle, connective tissue, goblet cells, ionocytes, glial cells, platelets, erythrocytes, endothelial cells, various immune cell types, and epidermal cells of multiple types (Fig. 1e-g, Supplementary Fig. 1b, and Supplementary Fig. 2).

To assess changes to cell populations and gene expression patterns, we evaluated cell clusters across regeneration stages. At 1 dpa, the major changes observed were the increases in proliferating cells, myeloid cells, lymphoid cells, and wound epidermis (Fig. 2a, b, and e), consistent with wound closure completion and immune cell infiltration observed via histology (Fig. 1d). At 3 dpa, all immune cell types were increased relative to 1 dpa, an indicative of substantial immune cell infiltration, yet the most notable change was the dramatic increase in erythrocytes (Fig. 2c and e). Massive immune cell infiltration, especially macrophages, and the formation of erythrocytes clumps in the limb blastema, have both been identified as features of salamander limb regeneration^15,16^. In the Japanese fire-bellied newt (*Cynops pyrrhogaster*), erythrocyte clumps express genes encoding secreted growth factors and matrix metalloproteases, which may play a role in limb regeneration^16^. In contrast, our analysis of differentially expressed genes in the erythrocyte cluster versus all other clusters found no strong evidence supporting *Polypterus* erythrocytes as a significant source of secreted growth factors or extracellular matrix remodelers (Supplementary Table 1). Finally, we observed reduction of muscle, connective tissue, ionocytes, goblet cells, and suprabasal epidermis clusters at 3 dpa, and a reversal of this trend at 7 dpa (Fig. 2c, d and e).

**Fig. 2.**
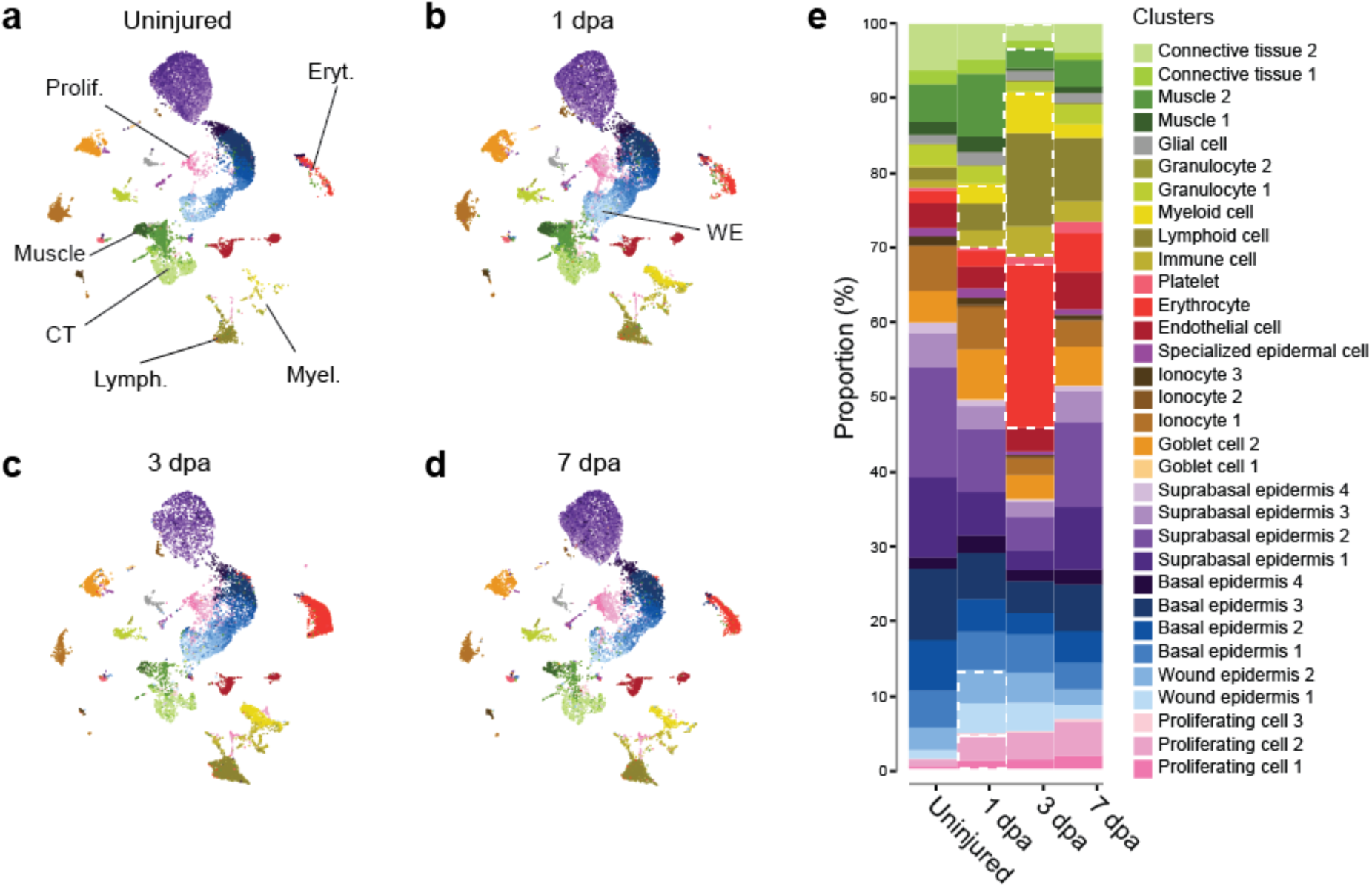
Fin regeneration is marked by reduction in muscle tissue and increased cell proliferation, immune cell and erythrocyte infiltration. UMAP of cell clusters at the uninjured stage (**a**) and across fin regeneration stages 1 dpa (**b**), 3 dpa (**c**) and 7 dpa (**d**); most dynamic cell populations were proliferative cells (Prolif.), erythrocytes (Eryt.), muscle, connective tissue (CT), lymphoid cells (Lymph.), myeloid cells (Myel.) and wound epidermis (WE). **e** Frequency plot showing proportions of cell clusters across regeneration stages; dashed white boxes highlight most dynamic clusters.

### Cellular contributions to the *Polypterus* fin wound epidermis and blastema

The formation of a signaling-competent wound epidermis and the recruitment of blastemal progenitor cells are key drivers of appendage epimorphic regeneration. Here we assessed the cellular contributions and gene expression changes in these cell populations from homeostasis and across regeneration stages. Our snRNA-seq unbiased clustering identified *col17a1b* and *col6a3* as markers for wound epidermis clusters, and for connective tissue clusters, respectively (Fig. 1f, g). In the uninjured fin, *col17a1b*-expressing cells were identified and grouped with the cluster designated as wound epidermis (Fig. 3a). This cell population expanded at 1 dpa (Fig. 2e, Fig. 3b), gradually declined by 3 dpa (Fig. 2e, Fig. 3c), and ultimately returned to a proportion similar to that of the uninjured fin by 7 dpa (Fig. 2e, Fig. 3d). Further analysis revealed that these wound epidermis-like cells in the uninjured fin likely represent a subpopulation of basal epidermal cells present during homeostasis. These cells share key markers with the wound epidermis clusters observed during regenerative stages, including *col17a1b* and *itga6a* (Supplementary Fig. 3). Connective tissue cells expressing *col6a3* were present during homeostasis and across all stages of regeneration, peaking in abundance at 1 dpa, similar to the wound epidermis (Fig. 2e, Fig. 3a-d). To confirm the identity of wound epidermis cells and the location of connective tissue cells at homeostasis and during regeneration, we used hybridization chain reaction (HCR) fluorescence *in situ* hybridization (HCR-FISH) staining. In the uninjured fin (Fig. 3e), *col17a1b*-expressing cells were detected, as anticipated, in cells along the basal layer of the wound epidermis. Presumptive connective tissue cells expressing *col6a3* were found in the dermis and dispersed between muscle cells. At 1 dpa (Fig. 3f), *col17a1b*-expressing cells were also detected in the basal layer of the wound epidermis, and clusters of *col6a3*-expressing cells were observed in the stump, near the amputation site. At 3 dpa (Fig. 3g), *col6a3*-expressing cells accumulated distally in the regenerating fin and were abundant immediately subjacent to *col17a1b*-expressing cells, in the presumptive blastema. At 7 dpa (Fig. 3h), *col6a3*-expressing cells were most abundant distally, and *col6a3* signal intensity was higher in cells closest to the *col17a1b*-expresing cells in the wound epidermis basal layer. These data indicate that *col17a1b* is a marker of a subset of basal epidermis cells at homeostasis and during regeneration, suggesting that *col6a3*-positive connective tissue cells are major contributors to the *Polypterus* fin blastema.

**Fig. 3.**
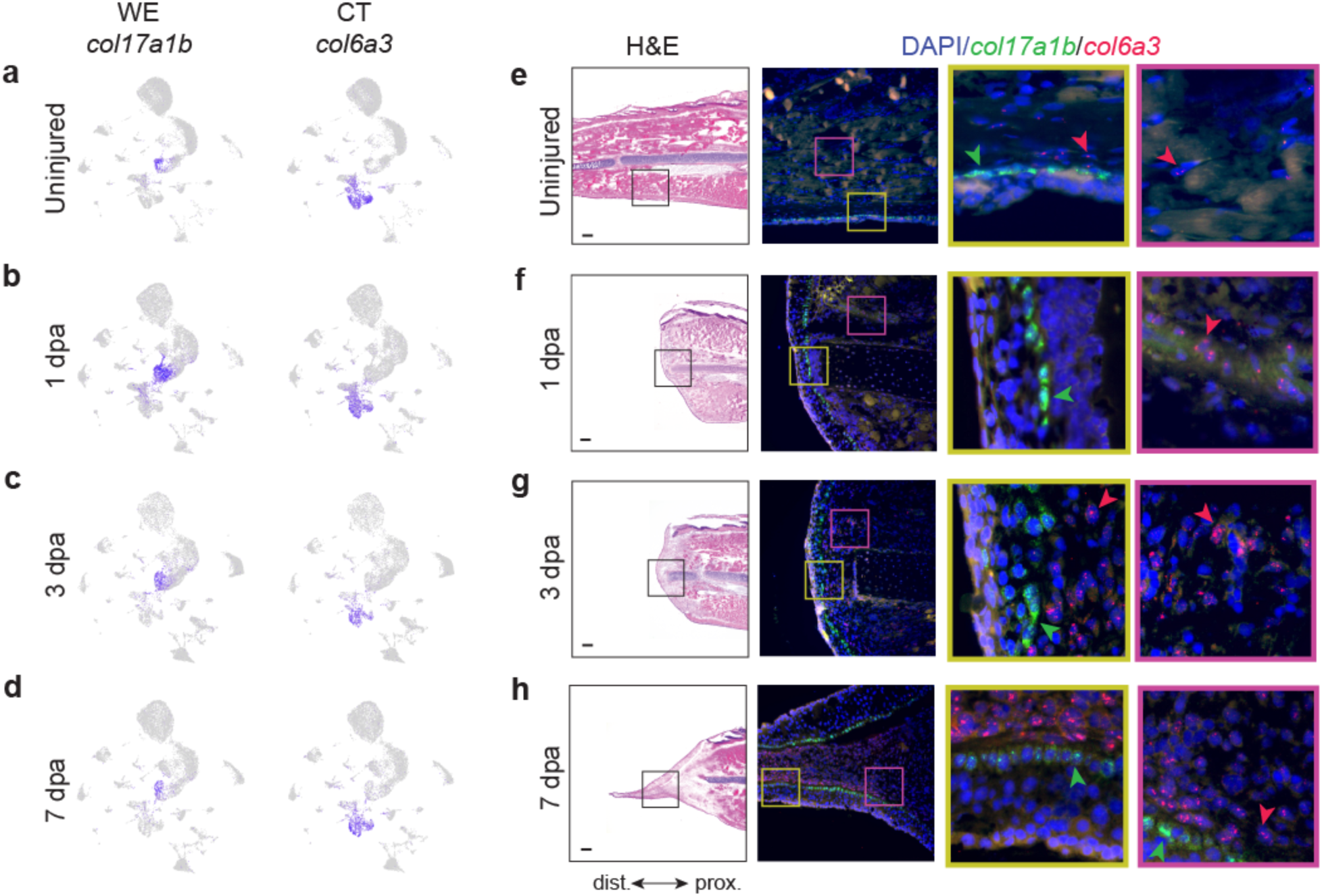
Cellular contributions to the fin wound epidermis and blastema. UMAP of cell clusters showing *col17a1b* and *col6a3* gene expression at the uninjured stage (**a**) and across fin regeneration stages 1 dpa (**b**), 3 dpa (**c**), and 7 dpa (**d**). Histology and HCR-FISH at the uninjured stage (**e**) and across fin regeneration stages 1 dpa (**f**), 3 dpa (**g**), and 7 dpa (**h**); HCR-FISH panels show DAPI staining (blue), *col17a1b* (green) and *col6a3* (red) gene expression; black box in the histology panels corresponds to approximate region in adjacent HCR-FISH panels; red and yellow boxes denote zoomed in regions showing higher magnification panels of HCR-FISH signal; green and red arrowheads denote *col17a1b*- and *col6a3*-expressing cells, respectively. WE, wound epidermis; CT, connective tissue; H&E, hematoxylin and eosin. Scale bars = 100 μm.

### SnRNA-seq and spatial RNA-seq reveal blastema cell heterogeneity and genetic markers of the wound epidermis

To identify the gene expression territories of wound epidermis and blastema markers, we generated high throughput, spatially resolved gene expression data using Visium spatial transcriptomics technology (10x Genomics). To compare spatial gene expression profiles between the *Polypterus* fin and the axolotl limb, we obtained histological sections of blastemas from both species at 3 dpa, a stage in which a multilayered wound epidermis and blastema cells are present in both species^3,4^ (Fig. 4a, e). Unbiased *k*-clustering analysis (*k* = 8) using Loupe Browser v8.0.0 (10x Genomics) readily identified major tissue types (Fig. 4b, c, f, g) and orthologous gene markers for blastema (*tnc*), wound epidermis (*palld*), muscle (*myoz1*), and skeleton (*col9a1*), in comparable regions of the *Polypterus* fin and axolotl limb at 3 dpa (Fig. 4d, h).

**Fig. 4.**
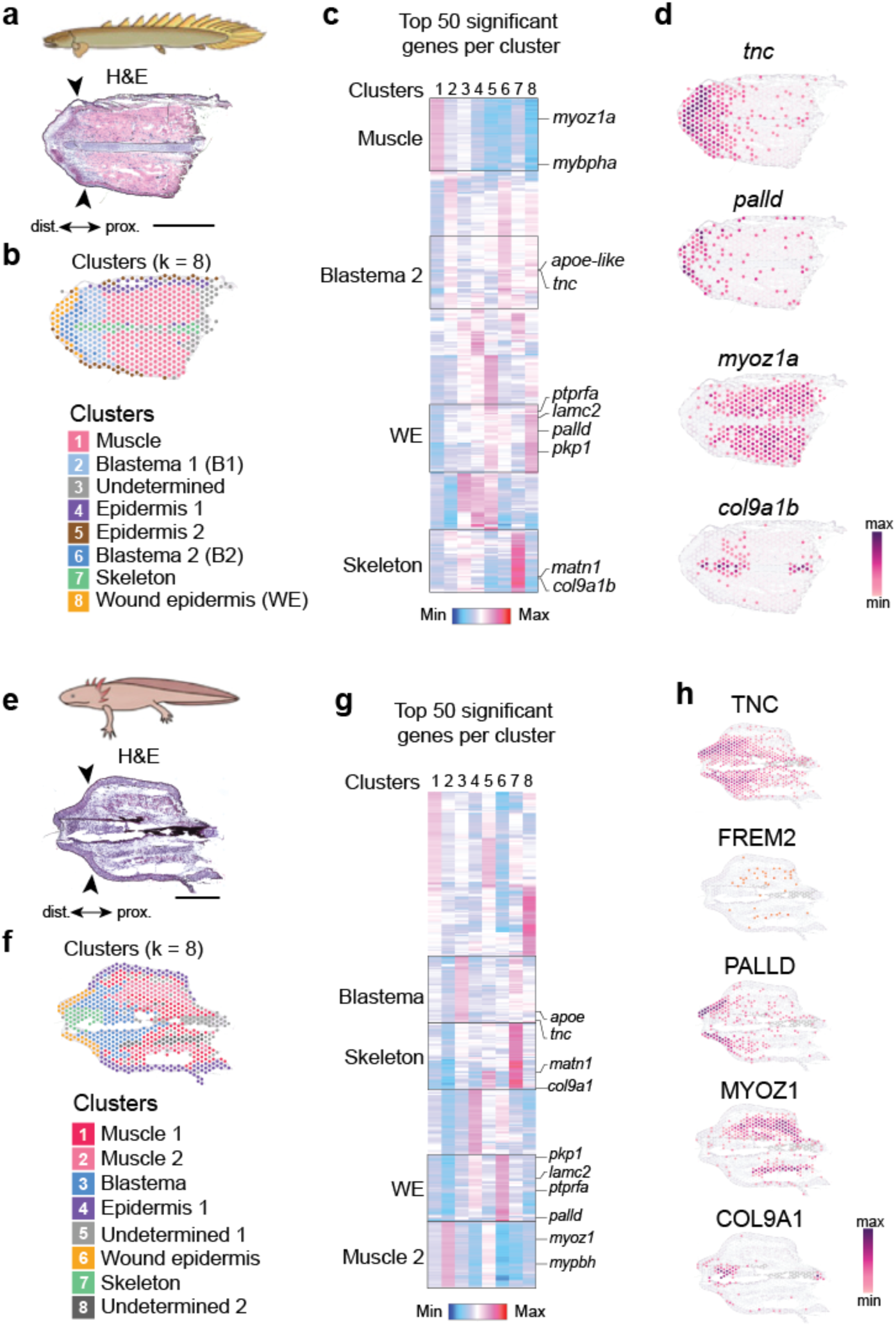
Spatial RNA-seq identifies conserved domains with distinct genetic signatures, underscoring spatial similarities of limbs and fins at 3 dpa. Histological sections used for spatial RNA-seq of *Polypterus* fin (**a**) and axolotl limb (**e**) at 3 dpa; black arrowheads indicate amputation plane. Unbiased k-means clusters identify distinct gene expression territories corresponding to muscle, skeleton, epidermis, wound epidermis, and blastema in the *Polypterus* fin (**b**) and axolotl limb (**f**) at 3 dpa. Heatmap showing top 50 significant genes per cluster at 3 dpa in *Polypterus* fin (**c**) and axolotl limb (**g**). Spatial expression profiles *Polypterus* fin genes (**d**) and their axolotl orthologs (**h**) show equivalent domains corresponding to the blastema (*tnc*), wound epidermis (*palld*), muscle (*myoz1a*), and skeleton (*col9a1b*); axolotl FREM2 expression at 3 dpa is mostly proximal to the amputation site.

Next, we examined more closely the identity of our snRNA-seq clusters annotated as wound epidermis and connective tissue cells by assessing the expression profiles of established axolotl limb regeneration markers across *Polypterus* fin regeneration stages. Expression of *mmp19*, a marker of axolotl limb blastema cells^17,18^ was found in a subset of the *Polypterus* fin connective tissue cells (Fig. 5a). Although seemingly dispensable for axolotl limb regeneration^19^, PAX7+ muscle satellite cells contribute to the axolotl limb blastema^20^. In the uninjured *Polypterus* fin, *pax7*a-expressing cells were present in the muscle cluster. Later, at 1 dpa and at 3 dpa, *pax7*a-expressing cells were also detected within the connective tissue clusters (Fig. 5b). Previous studies identified FREM2 as a marker of wound epidermis cells in the axolotl limb^21,22^. In *Polypterus*, *frem3*, not *frem2*, was expressed in a distinct subset of wound epidermis cells (Fig. 5c). MMP13 is highly expressed in the axolotl limb wound epidermis^22,23^. Likewise, the *Polypterus* ortholog, *mmp13b*, was highly expressed in the wound epidermis clusters (Fig. 5d). Interestingly, *kazald2*, which is highly expressed in the axolotl limb blastema and required for proper limb regeneration^24^, was expressed almost exclusively in the *Polypterus* wound epidermis clusters at 3 dpa and 7 dpa (Fig. 5e). This pattern mirrored that of *frem3*, as well as *and1* (Fig. 5f), a gene exclusively present in fish, which encodes actinodin 1, an essential structural component of actinotrichia^25^.

**Fig. 5.**
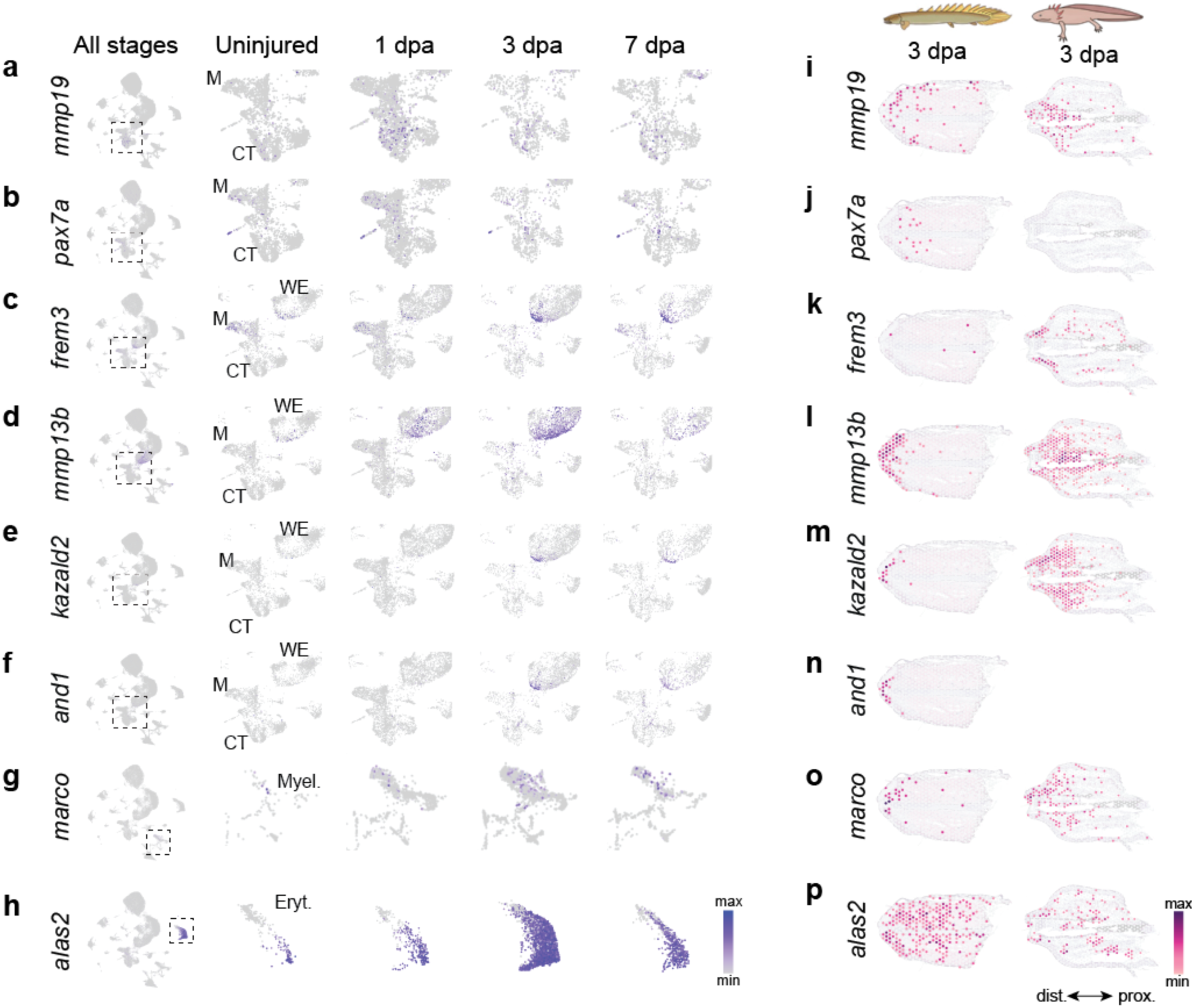
Spatial and snRNA-seq identify shared and derived wound epidermis markers, and heterogenous cellular contributions to the fin blastema. UMAP showing expression of *mmp19* (**a**), *pax7a* (**b**), *frem3* (**c**), *mmp13b* (**d**), *kazald2* (**e**), *and1* (**f**), *marco* (**g**), and *alas2* (**h**) in all stages combined, uninjured, 1 dpa, 3 dpa, and 7 dpa. Box with dashed lines in UMAP denotes zoomed in region in subsequent panels. Spatial RNA-seq expression profiles of *mmp19* (**i**), *pax7a* (**j**), *frem3* (**k**), *mmp13b* (**l**), *kazald2* (**m**), *and1* (**n**), *marco* (**o**), and *alas2* (**p**) in the *Polypterus* fin and axolotl limb at 3 dpa. M, muscle; CT, connective tissue; WE, wound epidermis; Myel., myeloid cell; Eryt., erythrocyte.

We next assessed the spatial expression of these markers in *Polypterus* and axolotl wound epidermis and blastema. Expression of *mmp19*, which was detected in a subset of connective tissue cells via snRNA-seq, was found mostly in the blastema region, subjacent to the wound epidermis, in both *Polypterus* and axolotl (Fig. 5i). Expression of *pax7a* was also located in the presumed blastema of the *Polypterus* fin at 3 dpa (Fig. 5j), yet no expression was detected in the axolotl limb, possibly due to the limited sensitivity of spatial RNA-seq^26^. Only a few spots positive for *frem3* were detected, either in the wound epidermis or scattered in the fin stump (Fig. 5k). In the axolotl, FREM2 expression was detected in the limb stump at 3 dpa (Fig. 4h), whereas FREM3 was conspicuously expressed in the wound epidermis, in addition to scattered spots in the fin stump (Fig. 5k). Both the *Polypterus mmp13b* and the axolotl MMP13 were detected in the wound epidermis and in immediately subjacent spots (Fig. 5l). The axolotl MMP13 was broadly expressed, possibly due to a larger presumptive blastema cell population in the axolotl limb compared to the *Polypterus* fin at this stage, as suggested by the expression patterns of MMP19 and TNC (Fig. 5i and Fig. 4h). As suggested by our snRNA-seq results, spatial RNA-seq confirmed distal expression of *kazald2* (Fig. 5m) and *and1* (Fig. 5n) in a subset of cells in the *Polypterus* fin wound epidermis. Conversely, KAZALD2 was found in both the axolotl limb wound epidermis and blastema (Fig. 5m).

Myeloid cells, particularly macrophages, infiltrate the limb blastema during regeneration. Our snRNA-seq dataset identified a substantial increase in the proportion of myeloid cells during regeneration, relative to homeostasis (Fig. 2a-e). To assess whether these cells reach the fin blastema, we examined the expression of *marco*, which marks a subpopulation of myeloid cells in the axolotl limb blastema^21,27^. As seen in the axolotl, our snRNA-seq dataset showed *marco*-expressing cells in the myeloid cluster during *Polypterus* fin regeneration (Fig. 5g). Spatial RNA-seq showed expression of *marco* in the presumptive blastema of the *Polypterus* fin and axolotl limb (Fig. 5o). Finally, *alas2* expression specifically marked a population of erythrocytes that increased massively during fin regeneration (Fig. 5h). Spatial RNA-seq showed *alas2* expressed in the blastema region and in spots located more proximally, in the fin stump (Fig. 5p). In the axolotl limb, ALAS2 expression was detected in clusters of spots along the limb, both in the blastema region and in proximal regions of the limb stump (Fig. 5p). Altogether, these findings confirm *frem3*, *mmp13b*, *kazald2* and *and1* as markers of *Polypterus* fin wound epidermis and suggest that, as seen in axolotls, the *Polypterus* fin blastema is composed of a heterogeneous cell population, with contributions from tissues such as presumptive muscle satellite cells (*pax7a*), connective tissue cells (*mmp19*), myeloid cells (*marco*) and erythrocytes (*alas2*).

The combination of snRNA-seq and spatial RNA-seq data also allowed us to better refine cell cluster annotation and identify subsets of cell types within clusters. In the specialized epidermal cell (SEC) cluster, the expression of *pou2f3* and *trpm5*, and the epidermal expression of *pou2f3* revealed by spatial RNA-seq, suggest that SEC may correspond to or contain tuft cells, an epithelial cell type involved in immune response, normally found in intestinal epithelium of tetrapods and in the skin of fish^28,29^ (Supplementary Fig. 4a). Some cells within the loosely annotated immune cluster showed expression of *syk* and *pax5*, both associated with B-cell identity^30,31^, suggesting that B-cells comprise at least a subset of this cluster (Supplementary Fig. 4b). Finally, skeletal cells were found to correspond to a subset of connective tissue cells, as evidenced by the expression of the markers *col2a1* and *matn1*, detected via spatial RNA-seq in the *Polypterus* fin skeleton (Supplementary Fig. 4c).

### Hypoxia response mechanisms during *Polypterus* fin regeneration

Evidence from various regenerative models including mouse^32^, *Xenopus*^33^, axolotl^34^ and zebrafish^35^, support a scenario where wounding drives local reactive oxygen species (ROS) production, which in turn promotes a hypoxic environment permissive to regeneration. The hypoxia-inducible factor 1 alpha (HIF1A) plays a crucial role in translating changes of oxygen levels into cellular responses essential for wound healing and regeneration^36^. HIF1A levels are tightly regulated by prolyl hydroxylase domain (PHD) enzymes (encoded by the EGLN1, EGLN2 and EGLN3 genes) and by the von Hippel–Lindau protein (pVHL, encoded by the VHL gene), which promote HIF1A degradation, and HIF1AN, (encoded by the HIF1AN gene), which negatively regulates HIF1A transcription. Hypoxia prevents degradation of HIF1A by its negative regulators, resulting in HIF1A-induced increase in expression of glycolytic enzymes^37,38^.

Recent findings demonstrated that during *Xenopus* and axolotl limb regeneration, the expression of HIF1A negative regulators is low, while HIF1A gene and glycolysis-related genes are highly expressed^39^. This suggests that these species stabilize HIF1A by maintaining low expression levels of its regulators during regeneration. We examined the expression of the *Polypterus* HIF1A ortholog, *hif1ab*, and its regulators, as well as glycolysis genes, in our spatial and snRNA-seq datasets. As seen in *Xenopus* and axolotl, *Polypterus egln1, egln2, egln3* and *hif1an* expression levels remain mostly low at both the uninjured fin and at 1 dpa, whereas *hif1ab* expression is greatly increased (Fig. 6a). At 3 dpa, even as expression of HIF1A regulators increase and *hif1ab* expression decreases (Fig. 6a), *hif1ab* remains broadly expressed in the blastema region, in contrast to its regulators (Fig. 6b). Concordantly, expression dynamics of glycolysis genes largely mirror that of *hif1ab* (Fig. 6a), and most are broadly expressed in the *Polypterus* fin at 3 dpa, especially in the muscle (Fig. 6c). Altogether, these findings suggest an evolutionarily conserved regenerative response to hypoxia, involving HIF1A expression stabilization and activation of glycolysis.

**Fig. 6.**
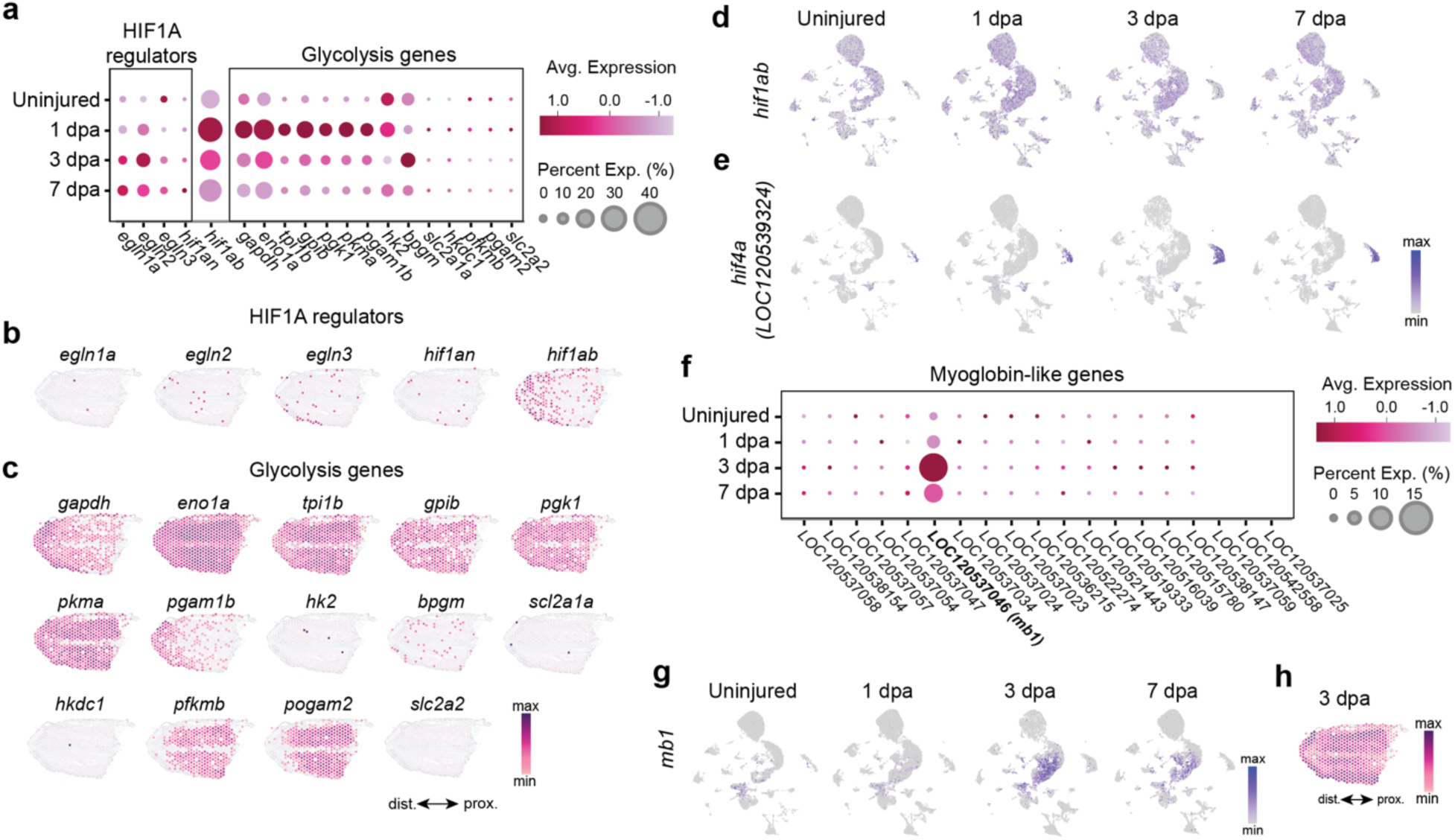
Hypoxia responses during *Polypterus* fin regeneration involve regulation of HIF1A, recruitment of *hif4a*+ erythrocytes and upregulation of myoglobin gene expression. **a** Dot plot showing expression of HIF1A regulators and glycolysis genes during fin regeneration. Spatial expression patterns of *hi1ab*, its regulators (**b**) and glycolysis genes (**c**) at 3 dpa. UMAP showing expression of *hif1ab* (**d**) and *hif4a* (LOC120539324) (**e**) in the uninjured fin and during regeneration stages. **f** Dot plot of myoglobin-like gene expression during fin regeneration. **g** UMAP showing expression of the *mb1* (LOC120537046) in the uninjured fin and during regeneration. h *mb1* spatial expression profile at 3 dpa.

In addition to the conserved hypoxia-HIF1A-glycolysis axis, our analyses also pointed to the involvement of genes in fin regeneration that are absent from tetrapod genomes. Whereas snRNA-seq shows that *hif1ab* upregulation occurs across all cell clusters (Fig. 6d), we detected the upregulation of *hif1a-like* (LOC120539324) almost exclusively in the erythrocyte cluster, which greatly increases during regeneration (Fig. 6e). Based on recent phylogenetically study of HIFA gene evolution in vertebrates^40^, we inferred that *hif1a-like* corresponds to the *hif4a* gene, a homolog to the ancestral vertebrate HIFA gene, which was retained in only some ray-finned fish lineages, including zebrafish.

Myoglobin (MB) is a versatile protein shown to mitigate both ROS activity and hypoxia. Specifically, high MB levels in muscle cells can increase oxygen storage capacity in marine mammals during extended dives^41^. Furthermore, MB attenuates oxidative stress in cardiac muscle^42^. Remarkably, while most vertebrates possess a single MB gene, *Polypterus* possesses at least 15 *mb* genes^43^, whereas frogs and salamanders have lost their MB orthologs^44^. We examined the expression of *Polypterus mb* genes in our snRNA-seq dataset and found 19 genes currently annotated as myoglobin-like genes in the *Polypterus* genome. Of those, LOC120537046 was markedly upregulated during fin regeneration (Fig. 6f). A previous study showed that among *Polypterus* myoglobin-like proteins, PseMb1 the product of LOC120537046, termed PseMb1, showed the highest identity to the mammalian MB gene^43^. Here, we refer LOC120537046 as *mb1*. In the uninjured and 1 dpa fin tissue, *mb1* was expressed in the muscle and in a few cells in the basal epidermis clusters (Fig. 6g). At 3 dpa, *mb1* expression in the wound epidermis increased, and then moderately decreased at 7 dpa (Fig. 6g). Spatial RNA-seq showed *mb1* broadly expressed at 3 dpa, with the highest levels in the muscle and wound epidermis (Fig. 6h). While the roles of the *Polypterus hif4a* and *mb1* genes in fin regeneration remain unclear, these findings suggest species-specific mechanisms for adapting to oxygen stress, providing a foundation for further investigation into their functional significance.

### Activation of DNA damage response in cycling cells during fin regeneration

Previous studies on diverse regenerative models, including planarians^45^, newts^46^, *Polypterus,* and axolotl^4^, have demonstrated the upregulation of DNA repair genes at the onset of regeneration. Activation of DNA damage response (DDR) has been shown to be essential for regeneration of both the axolotl limb^47^ and the zebrafish heart^48^. Specifically, in the zebrafish heart, DDR is activated in cardiomyocytes near the injury site, particularly in those undergoing cell division^48^. In the axolotl limb, the genetic deletion of EYA2, a key regulator of H2AX phosphorylation—a major DNA damage mediator—preferentially affects cells that have entered the cell cycle^47^. Likewise, analysis of our snRNA-seq data revealed upregulation of DDR markers during fin regeneration. At 1 dpa, expression of the DNA damage sensing markers *atm*, *atr*, *tp53* and, to a lesser extent, *eya2*, was observed in a larger number of cells when compared to DNA damage repair markers *rad51*, *rpa2*, *fen1*, *chaf1a*, and *asf1bb* (Fig. 7a, Supplementary Fig. 5). Expression of *atm*, *atr* and *tp53* was found in every snRNA-seq cluster, including proliferating cells, marked by *mki67* expression (Fig. 7b, Supplementary Fig. 5). Expression of *eya2* was also detected in several clusters, including proliferating cells, but was more prevalent in cells of the basal epidermis and connective tissue clusters. Interestingly, expression of DNA damage repair markers was preferentially found in the proliferating cell clusters (Fig. 7a, e, f, Supplementary Fig. 5). Interestingly, the spatial distribution of proliferating cells and DDR markers was strikingly different between *Polypterus* and axolotl. In the *Polypterus* fin at 3 dpa, *mki67* and all analyzed DDR markers were expressed mostly distally, in the wound epidermis and blastema regions (Fig. 7c, f). Conversely, the axolotl orthologs were broadly expressed, indicating activation of cell proliferation and DDR in the wound site but also proximally, along the limb stump (Fig. 7d, g). Together, these findings suggest a common activation of DDR in cycling cells during the regeneration of the *Polypterus* fin and the axolotl limb. However, they also reveal an unexpected difference in the spatial distribution of these cells relative to the wound site between *Polypterus* and axolotl.

**Fig. 7.**
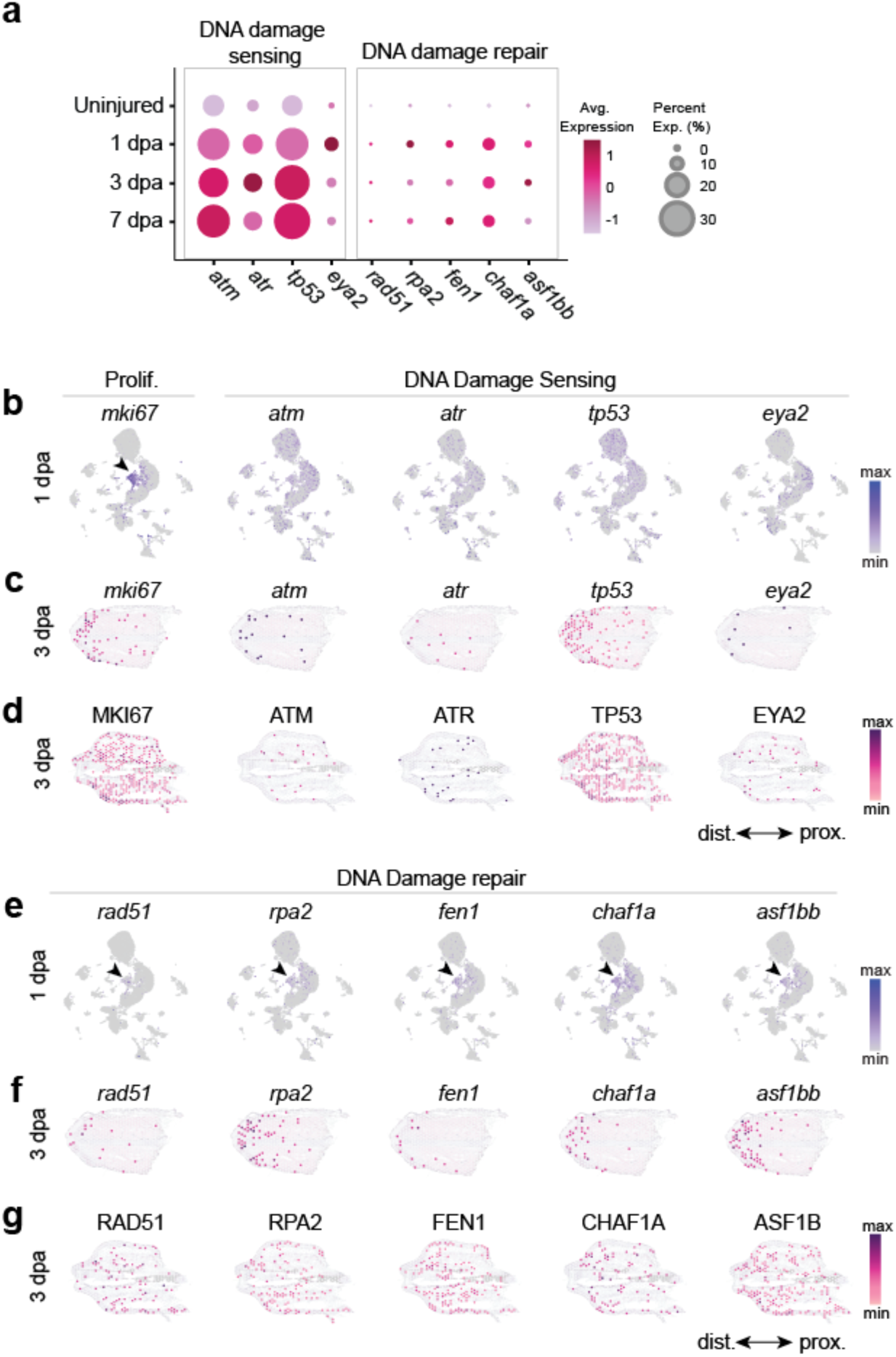
*Polypterus* fin amputation triggers DNA damage response in cycling cells near the injury site. **a** Dot plot showing expression of DNA damage sensing genes and DNA repair genes. SnRNA-seq UMAP (**b**), spatial RNA-seq of *Polypterus* fin (**c**) and axolotl limb (**d**), showing expression of cell proliferation and DNA damage sensing markers. SnRNA-seq UMAP (**e**), spatial RNA-seq of *Polypterus* fin (**f**) and axolotl limb (**g**), showing expression of DNA damage repair markers. Arrowheads indicate snRNA-seq cell proliferation clusters.

### Switch from pro-to anti-inflammatory immune response

Following axolotl limb amputation, pro- and anti-inflammatory signals are induced and sustained through regeneration^15^, with pro-inflammatory macrophages peaking during early to mid-blastema stages and anti-inflammatory macrophages being predominant at later stages^49^. Likewise, in larval zebrafish, fin amputation results in accumulation of pro-inflammatory macrophages at the wound site at 1 dpa, followed by an increase in anti-inflammatory macrophages^50^. Our analysis of the immune response landscape of *Polypterus* fin regeneration broadly parallels the pattern described in both the axolotl and zebrafish. Upon fin amputation, at 1 dpa, we observed upregulation *of il1b* expression, a classic marker of pro-inflammatory macrophages^51^, in a subset of myeloid cells (Fig. 8a,b). At 1 dpa, the myeloid cluster also expressed high levels of *spp1,* a fibrogenic macrophage marker^52^, which was sharply downregulated at subsequent stages. Other candidate orthologs of mammalian genes associated with pro-inflammatory macrophages, such as *tnfsf15*^53^ and *hsp90b1*^54^, were upregulated at 1 dpa. Anti-inflammatory markers were also upregulated at 1 dpa, but their expression peaked at subsequent regeneration stages (Fig. 8a, c). *tgfb1*, an established immune modulator frequently presenting anti-inflammatory activity, was gradually upregulated (Fig. 8a) and broadly expressed in cells of the myeloid cluster (Fig. 8c) during regeneration stages. Expression of other putative orthologs of anti-inflammatory markers, *hs3st1-like*, *clec79-like*, and *mcf2la*, also peaked at later stages of fin regeneration. While some overlap exists between pro- and anti-inflammatory marker expression at 1 dpa, differences in the subset of cells expressing *il1b*/*spp1* and *hs3st1l1*/*clec7a-like* within the myeloid cluster (Fig. 8b, c) suggest that pro- and anti-inflammatory macrophages coexist in early regeneration stages. These data support a similar dynamic of pro- and anti-inflammatory immune cell response during fin and limb regeneration.

**Fig. 8.**
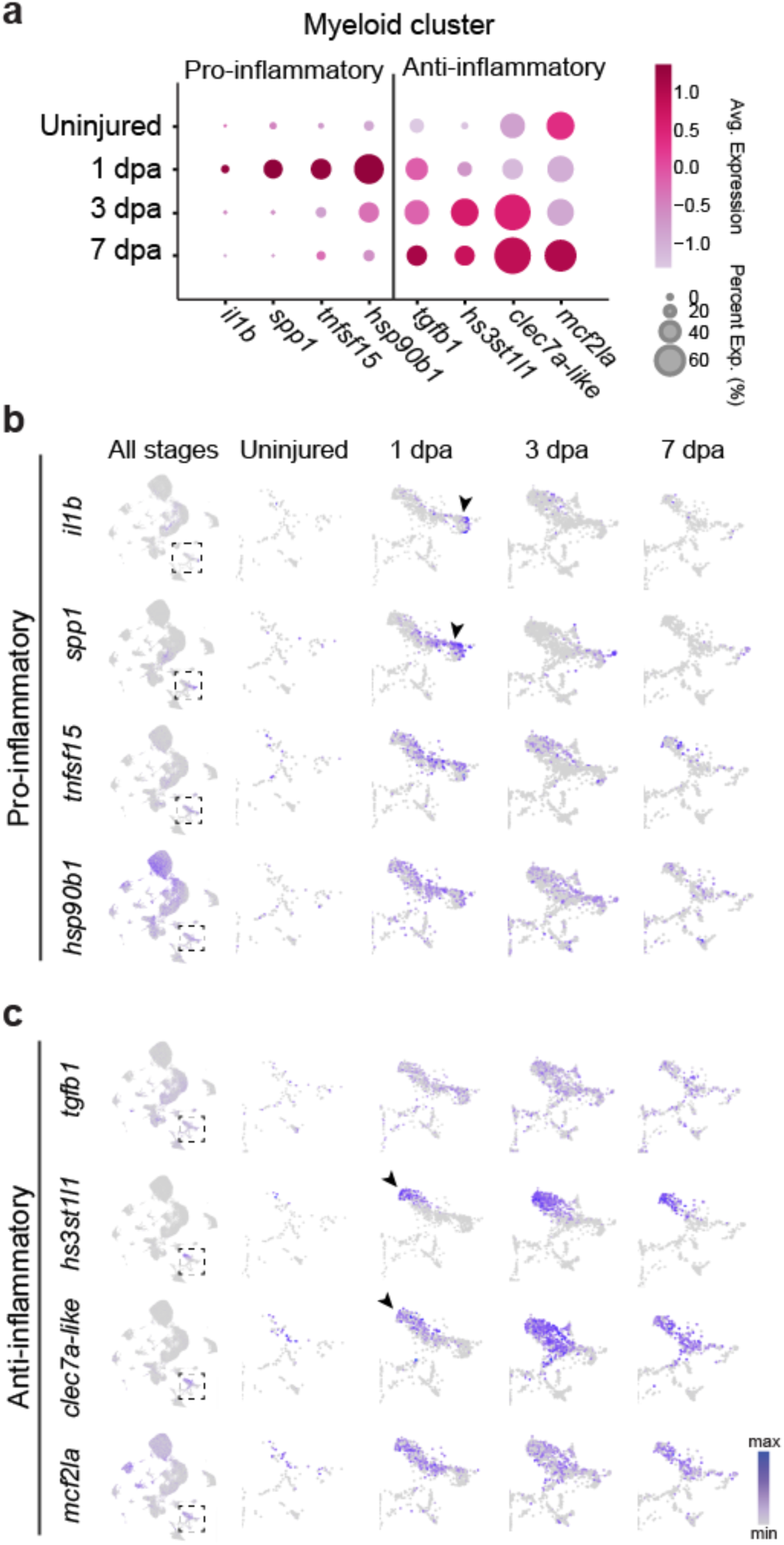
Switch from anti- to pro-inflammatory myeloid cell phenotype during *Polypterus* fin regeneration. **a** Dot plot showing expression of pro- and anti-inflammatory markers in the myeloid cell cluster in the uninjured fin tissue and across fin regeneration stages. SnRNa-seq UMAP expression of pro- (**b**) and anti-inflammatory (**c**) marker expression in the myeloid cell cluster. Box with dashed lines indicates location of myeloid cell cluster in the UMAP of all stages combined; subsequent panels show UMAP of myeloid cluster in the uninjured fin and regeneration stages. Arrowheads denote upregulation of marker genes in subsets of myeloid cells.

### Shared signaling factors expressed in the limb and fin wound epidermis and blastema

The comparative analysis of *Polypterus* fin and axolotl limb regeneration provides a unique opportunity to identify signaling components of an ancestral regenerative program for limbs and fins. To explore this, we examined the expression of genes encoding secreted and membrane-bound signaling factors in the wound epidermis and connective tissue clusters. Differential gene expression (DGE) analysis of snRNA-seq wound epidermis clusters at 1 dpa versus the uninjured fin uncovered several candidates, some with previously described roles in salamander limb regeneration, in including *marcksl1*^55^ (Fig. 9a, Supplementary Table 2). Testin, encoded by the gene *tes*, is a focal adhesion protein recently found to be involved cellular response to mechanical stimuli^56,57^. *tes* was robustly upregulated in the wound epidermis at 1 dpa and 3 dpa (Fig. 9b). Spatial RNA-seq showed *tes* expressed primarily in the *Polypterus* wound epidermis at 3 dpa, and in both wound epidermis and blastema in the axolotl limb (Fig. 9c). Another gene strongly upregulated in the wound epidermis was *tmem154*, which encodes a poorly characterized transmembrane protein previously implicated in the healing response to skin lesions caused by viral infections in sheep^58^. *S*nRNA-seq showed that, like *tes*, *tmem154* was upregulated at 1 dpa in the wound epidermis, with expression tapering off as regeneration progressed (Fig. 9b). At 3 dpa, spatial RNA-seq showed *tmem154* expression mostly restricted to the wound epidermis in both *Polypterus* and axolotl (Fig. 9c).

**Fig. 9.**
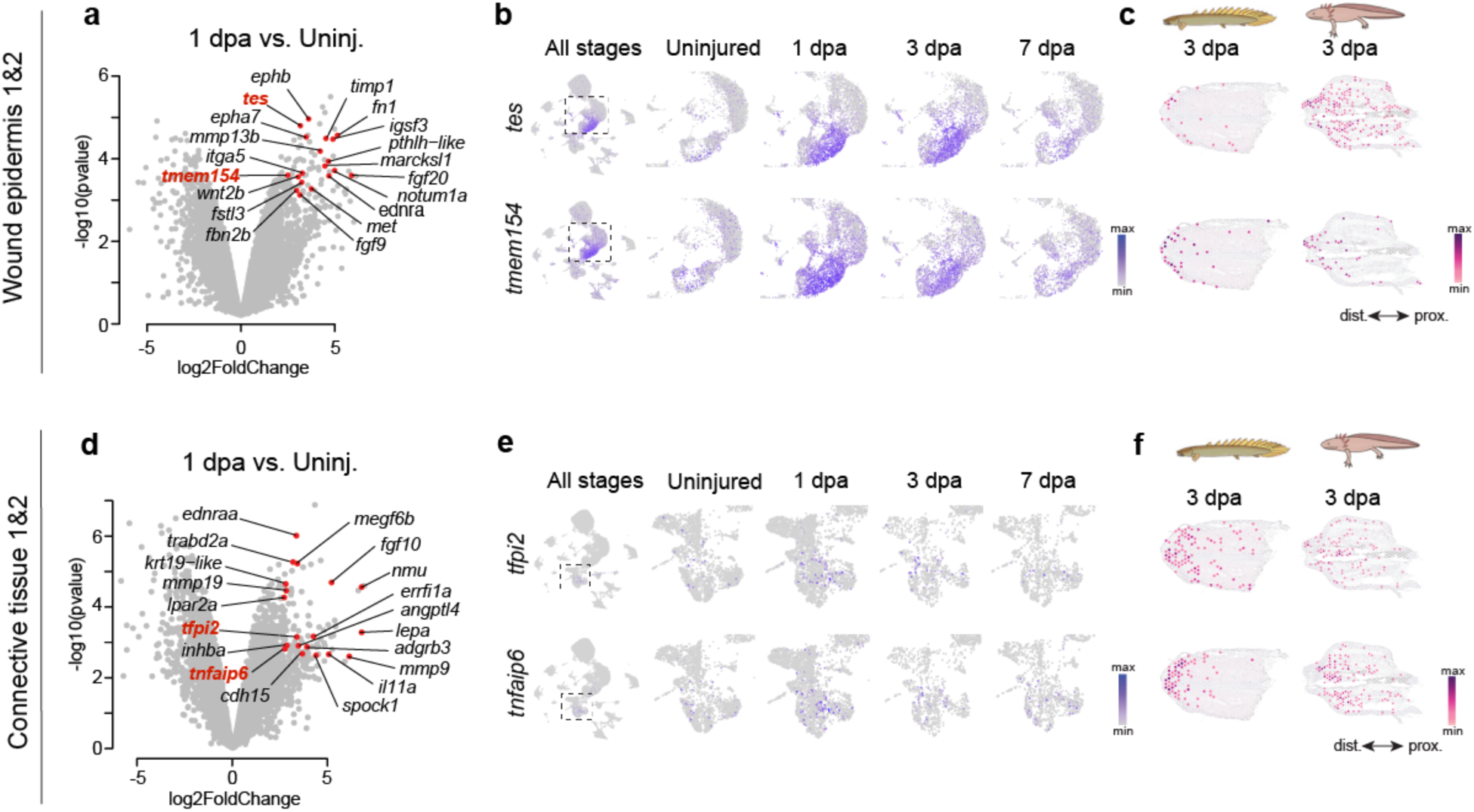
SnRNA-seq and spatial RNA-seq identify novel candidate signaling factors upregulated in the wound epidermis and blastema during fin and limb regeneration. **a** Volcano plot of genes upregulated in the *Polypterus* snRNA-seq wound epidermis clusters at 3 dpa relative to uninjured fin tissue that encode for membrane-bound or secreted proteins. **b** UMAP showing upregulation of *tes* and *tmem154* expression in the wound epidermis clusters. **c** Spatial expression of *tes* and *tmem154* in the *Polypterus* fin and axolotl limb at 3 dpa. **d** Volcano plot of genes upregulated in the *Polypterus* snRNA-seq connective tissue clusters at 3 dpa relative to uninjured fin tissue that encode for membrane-bound or secreted proteins. **e** UMAP showing upregulation of *tfpi2* and *tnfaip6* expression in the connective tissue clusters. **f** Spatial expression of *tfpi2* and *tnfaip6* in the *Polypterus* fin and axolotl limb at 3 dpa.

DGE analysis of connective tissue clusters at 1 dpa versus the uninjured fin revealed upregulation of genes encoding for established blastema signaling factors (Fig. 9d, Supplementary Table 3), such as inhibin beta A (*inhba*) and leptin (*lepa*), both upregulated in blastemas of the zebrafish fin ray^59,60^, the *Xenopus* tail^61,62^ and the axolotl limb^63,64^. Among the genes upregulated at 1 dpa was tissue factor pathway inhibitor 2, *tfpi2*, to our knowledge not previously implicated in limb or fin regeneration. *tfpi2* encodes a widely studied Kunitz-type protease inhibitor that has been associated with various processes, including regulation of tumor stemness^65^ and extracellular matrix remodeling during myocardial infarction^66^. We also detected upregulation of *tnfaip6*, which encodes TNF Alpha Induced Protein 6, and is required for proliferation and differentiation of mammalian mesenchymal stem cells^67^, and is upregulated in mesenchymal cells that will give rise to the zebrafish tail fin blastema^68^. SnRNA-seq showed upregulation of both *tfpi2* and *tnfaip6* in subsets of connective tissue cells (Fig. 9e). Spatial RNA-seq showed distal expression of both *tfpi2* and *tnfaip6*, largely overlapping with the blastema region (Fig. 9f). Together, these findings demonstrate the power of a comparative approach for the identification of novel candidate genes for downstream functional studies.

### Epigenetic analysis uncovers candidate fin regeneration-responsive elements

The identification of Tissue Regeneration Enhancer Elements (TREEs)^60^ active during limb and fin regeneration may ultimately reveal conserved gene regulatory networks that orchestrate a successful regenerative program. Various TREEs active during teleost fish fin ray regeneration have been identified in recent years^8,69–71^, yet changes to the epigenetic landscape in response to injury at the fin endoskeleton region, the homologous counterpart to the tetrapod limb, remain entirely unexplored. To address this, we used ATAC-seq to assess chromatin accessibility in the uninjured fin and at 3 dpa, a stage in which a signaling-competent wound epidermis is established, and blastemal cells are readily detected. Biological triplicates of uninjured and 3 dpa fin samples were used for ATAC-seq library preparation. Over 150 million paired-end reads were sequenced per sample, and the resulting libraries showed the expected fragment periodicity, separation between conditions, and clustering between replicates, along with over thirty-two and forty-seven thousand peaks called for the uninjured and 3 dpa conditions, respectively (Supplementary Fig. 6, Supplementary Table 4). Our analysis revealed 1,427 differentially accessible chromatin regions (adjusted p-value < 0.05), or peaks, between the 3 dpa and the uninjured fin tissue (Fig. 10a, Supplementary Table 5). Of those, 155 peaks mapped to unplaced DNA scaffolds. The 1,272 peaks mapped to *Polypterus* chromosomes were mostly (>86%) intergenic (Supplementary Fig. 6).

**Fig. 10.**
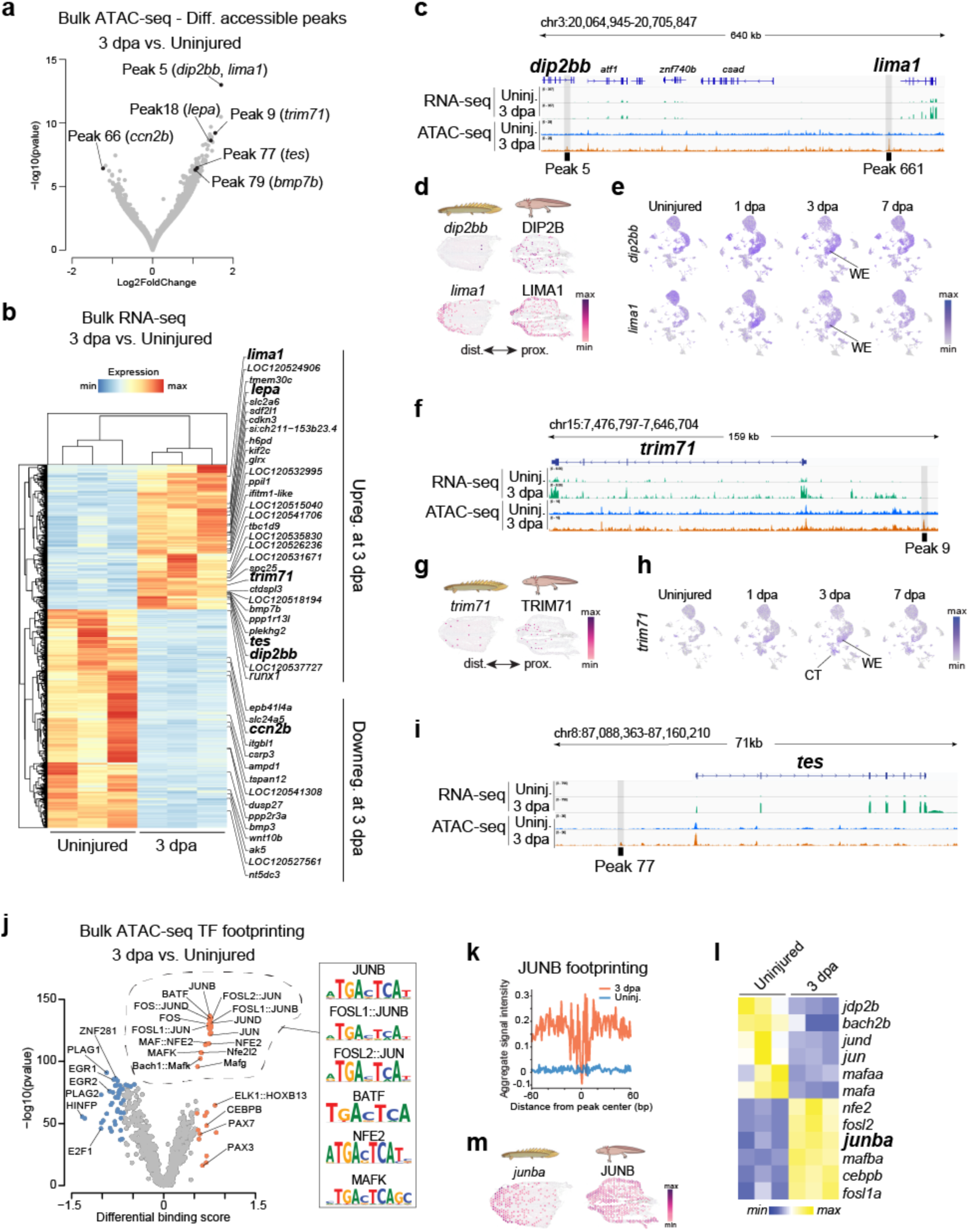
Epigenetic profiling of *Polypterus* fin regeneration uncovers candidate TREEs and enrichment of AP-1 TF footprint. **a** Volcano plot showing differentially accessible chromatin regions at 3 dpa relative to uninjured fin tissue. **b** Heatmap of bulk RNA-seq showing up and downregulated genes at 3 dpa relative to uninjured fin tissue. Genomic location of peaks 5 and 661 within *dip2bb* and upstream of *lima1* (**c**), peak 9 upstream of *trim71* (**f**), and peak 77 upstream of *tes* (**i**), including bulk RNA-seq and ATAC-seq tracks. Spatial expression of *Polypterus dip2bb* and *lima1* (**d**), *trim71* (**g**), and their respective axolotl orthologs at 3 dpa. SnRNA-seq UMAP showing expression of *dip2bb* and *lima1-like* (**e**) and *trim71* (**h**) in the uninjured fin and across regeneration stages. **j** Volcano plot of TF differential binding scores from ATAC-seq peaks at 3 dpa relative to uninjured fin tissue; dashed line denotes AP-1 TF family members and their preferred binding sites. **k** Aggregate plot across all JUNB binding sites. **l** AP-1 TFs differentially expressed in bulk RNA-seq data at 3 dpa relative to uninjured fin tissue. **m** Spatial expression of *Polypterus junba* and its axolotl ortholog. WE, wound epidermis. CT, connective tissue.

Focusing on the top 100 differentially accessible peaks, ranked by statistical significance, we searched for potential gene targets, reasoning that genes within their regulatory range would be differentially regulated during fin regeneration. To this end, we generated bulk RNA-seq data from uninjured and 3 dpa *Polypterus* fins in biological triplicates. Over 40 million paired-end reads were sequenced per sample (Supplementary Table 6), conditions showed proper separation and replicates clustered. We found 4,393 differentially expressed genes (Fig. 10b, Supplementary Table 7). Next, we searched for differentially expressed genes (fold change > 2, adjusted p-value < 0.05) within 600 kb up or downstream of our top 100 intergenic peaks. For peaks located in a gene desert, we expanded the search to 1.2 mb up or downstream of the peak. We found 46 genes upregulated at 3 dpa located in the vicinity of our top 100 ATAC-seq peaks (Fig. 10b). This list included genes implicated in limb regeneration from prior studies, such as *lepa* (Supplementary Fig. 7a-c), and *bmp7b* (Supplementary Fig. 7a-c). In addition, we detected *trim71*, recently shown to potentiate proliferation of reprogrammed limb progenitors^72^ and *tes*, identified in our snRNA-seq dataset as upregulated in the wound epidermis of both the *Polypterus* fin and the axolotl limb (Fig. 9a-c, Fig. 10b). Among the peaks differentially accessible in the uninjured fin, we found peak 66, located near *ccn2b*, a gene highly expressed at homeostasis and downregulated over 6-fold at 3 dpa (Fig. 10a,b, Supplementary Fig. 7g,h, Supplementary Table 7). Its downregulation at 3 dpa is consistent with its reported role in promoting fibrosis^73^.

Next, we inspected more closely 3 peaks ranked among the top 20 most significantly changed from the uninjured to the 3 dpa fin stage: peak 5 (within an intron of *dip2bb* and upstream of *lima1*), peak 9 (upstream of *trim71*), and peak 77 (upstream of *tes*). Peak 5, located in intron 22 of *dip2bb*, had two genes upregulated in its vicinity: *dip2bb* and *lima1* (Fig. 10c). While both genes are upregulated by just over 2-fold, bulk RNA-seq showed that *lima1* expression at 3 dpa is over 15 times greater than that of *dip2bb* (Supplementary Table 7). Of note, this genomic landscape also included peak 661, located 19 kb upstream of *lima1-like*. Consistent with the low expression level, Spatial RNA-seq detected only a few positive spots of *dip2bb* expression in the *Polypterus* fin stump at 3 dpa. Its axolotl ortholog, DIP2B, was detected in scattered spots in the epidermis and mesenchyme (Fig. 10d). SnRNA-seq showed increased *dip2bb* expression across all epidermal cell clusters, including the wound epidermis during regeneration stages (Fig. 10e). Differently, *lima1-like* and its axolotl ortholog LIMA1 were expressed throughout the epidermis, but noticeably highly expressed in the wound epidermis (Fig. 10d). Concordantly, snRNA-seq showed robust *lima1-like* expression in basal and suprabasal epidermal clusters, with increasing levels of expression in the wound epidermis as regeneration progressed (Fig. 10e). Peak 9 is located 50 kb upstream of *trim71*, which was upregulated at 3 dpa in our bulk RNA-seq dataset (Fig. 10f). Spatial RNA-seq showed *trim71* expressed in scattered spots along the proximal and distal mesenchyme, and wound epidermis of both the *Polypterus* fin and the axolotl limb at 3 dpa (Fig. 10g). SnRNA-seq showed increased *trim71* expression during fin regeneration, particularly in the connective tissue and wound epidermis clusters (Fig. 10h). Finally, peak 77 was located approximately 13 kb upstream of *tes* (Fig. 10i).

Finally, we investigated genome-wide transcription factor (TF) binding dynamics using TOBIAS^74^. We analyzed differential TF occupancy, or footprinting, across the 1,427 differentially accessible peaks identified between 3 dpa and uninjured fin tissue. The most striking trend in our differential TF binding analysis was the enrichment of bound sites for AP-1 family TFs at 3 dpa (Fig. 10j, Supplementary Table 8). Among these, JUNB exhibited the most statistically significant differential binding at 3 dpa compared to uninjured fin tissue. Consistently, the aggregated JUNB footprint plot for all 1,427 peaks displayed a well-defined JUNB footprint signal in regions differentially accessible at 3 dpa relative to the uninjured fin (Fig. 10k). We examined the expression of *Polypterus* AP-1 TFs in bulk RNA-seq data from 3 dpa versus uninjured fins and found the *Polypterus* JUNB ortholog, *junba*, significantly upregulated at 3 dpa (fold change = 3.3, adjusted p-value = 2.25E-12) (Fig. 10l, Supplementary Table 8). Spatial transcriptomics data showed *Polypterus junba* and axolotl JUNB widely expressed in epidermal and mesenchymal cells, with *Polypterus* showing a more distally (towards the injury site) concentrated expression of *junba* (Fig.10m). Overall, our findings indicate that, similar to axolotls and zebrafish, AP-1 TFs play a crucial role in regulating the activity of *Polypterus* candidate TREEs.

## Discussion

Highly regenerative vertebrates may achieve complete limb or fin regeneration by employing conserved genetic and cellular programs inherited from their last common ancestor. At the same time, they may also activate species-specific, derived programs that evolved later and may rely on novel genes. While species-specific programs are confined to the model organism being studied, ancestral regenerative programs are expected to be shared across multiple regenerative species. Therefore, comparative studies of limb and fin regeneration are essential for distinguishing between ancestral and derived mechanisms, ultimately accelerating the discovery of the core genetic and cellular processes that drive successful regeneration. By utilizing *Polypterus* as a model system, we expand upon insights gained from teleost models such as zebrafish, extending our understanding of fin regeneration to include the endoskeletal region—the homologous counterpart to the tetrapod limb.

Our comprehensive snRNA-seq profiling of the *Polypterus* uninjured and regenerating fin revealed the major changes in cellular composition across regeneration stages. Many of these changes parallel those observed during salamander limb regeneration. These include the reduction of the proportion of cells of the muscle cluster during regeneration, likely due to the extensive muscle histolysis that occurs prior to blastema formation, and increase in proliferating cells, and immune cell types, particularly myeloid cells (Fig. 2). Likewise, we detected multiple layers of epidermal cells, including a basal layer expressing *col17a1b* (Fig. 3), which has also been detected in the axolotl^21^. Furthermore, we found a massive increase in *alas2* -expressing erythrocytes that, based on *alas2* spatial expression at 3 dpa, contribute to the fin blastema (Fig. 5). Differently than erythrocyte clumps in newts^16^, we found no evidence in support of *Polypterus* erythrocytes as a significant source of secreted factors to the blastema. As a result, their potential role during fin regeneration remains unclear. Interestingly, *Polypterus hif4a* was markedly upregulated in *alas2*-expressing erythrocytes. *hif4a* is an ohnolog^75^ of *hif1a*, originated from the two rounds of WGD events that occurred at the base of vertebrate evolution. The *hif4a* gene was lost in the lineage leading to tetrapods, but retained in many bony fish species including *Polypterus* and zebrafish^40^. Therefore, *alas2*-expressing erythrocytes in *Polypterus* may play a role in oxygen sensing and hypoxia signaling during regeneration.

Integration of snRNA-seq expression data with spatial RNA-seq allowed us to approximate the location of specific cell populations in the *Polypterus* fin at 3 dpa (Fig. 4). Combined with our axolotl spatial RNA-seq dataset, we were able to identify similar and discordant gene expression patterns during fin and limb regeneration (Fig. 5). In the wound epidermis, we found shared markers between *Polypterus* and axolotl, such as *mmp13b*, *frem3* and *palld*. Expression of *kazald2* in the regenerating fin was restricted to the wound epidermis, differently than that of axolotls. Previous studies demonstrated that typical genetic programs of the apical ectodermal ridge (AER) – the distal signaling epithelium that helps drive limb outgrowth during embryogenesis – are deployed instead by mesenchymal cells in the axolotl limb during development^76^ and regeneration^77^. More recently, comparative analysis of *Xenopus* tadpoles and axolotls showed that transcriptional programs of the *Xenopus* limb wound epidermis were activated in both the wound epidermis and subjacent mesenchyme during axolotl limb regeneration^22^. We postulate that *kazald2* is a component of this genetic program, ancestrally activated only in the wound epidermis, as seen in frogs and *Polypterus*, later coopted by mesenchymal cells during salamander limb regeneration.

Contrary to mammals, the urodele amphibians such as newts can regrow severed limbs even at high environmental oxygen concentrations^78^. Recently, it has been proposed that regenerative models such as the frog tadpole and the axolotl cope with high oxygen concentration by maintaining low expression levels of HIFA regulators during regeneration^39^. This would enable stabilization of HIF1A and activation of glycolytic genes regardless of oxygen levels. Our results suggest that a similar mechanism may be taking place during Polypterus fin regeneration (Fig. 6). Our snRNA-seq data showed moderate upregulation of HIF1A modulators, whereas *hif1ab* expression was robustly increased, as were the expression of glycolysis genes. At 3 dpa, our spatial RNA-seq showed *hif1ab* broadly and robustly expressed in the fin blastema, whereas only scattered spots were positive for expression of HIF1A regulators. In sum, these findings suggest that a reduced oxygen sensitivity via HIF1A stabilization may be a vertebrate ancestral trait. In addition to the retention of *hif4a* in *Polypterus* and other fish species, *Polypterus* experienced a remarkable expansion in their myoglobin gene repertoire^43^, whereas salamander and frog genomes lost their *mb* gene ortholog^44^. We identified a *Polypterus* myoglobin-like gene, termed *mb1*, as highly expressed in the muscle but, surprisingly, also in the wound epidermis. Myoglobin gene expression in non-muscle tissues is rare but has been reported in the liver, gills and brain of the common carp (*cyprinius carpio*) as a potential adaptation to low oxygen environments^79^ and in human carcinoma cells, also for coping with hypoxic conditions associated with neoplastic growth^80^. While the role of *mb1* in the fin wound epidermis is unclear, our findings identify species-specific mechanisms that may be in place to manage oxygen and metabolic demands of regeneration.

Activation of DNA damage response is a hallmark of limb and fin regeneration. Recently, DDR has been detected in zebrafish cycling cardiomyocytes located near the injury site^48^. In the axolotl, DDR is also activated in cycling cells during regeneration^47^. Here we find that this is also true during *Polypterus* fin regeneration (Fig. 7). However, while proliferating cells expressing DDR-related genes in the *Polypterus* are mostly confined to the distal portion of the fin, these cells in the axolotl limb at 3 dpa are widely distributed, in both the blastema region and the limb stump. These observations suggest a broader cell cycle activation upon injury in the axolotl compared to the *Polypterus*. This is consistent with previous findings that showed how axolotl limb amputation leads to systemic cell cycle activation throughout the body^81^. Further studies in *Polypterus* and other regenerative species will help establish whether a systemic activation of cell cycle – and DDR – is unique to axolotls or is an evolutionarily shared feature of limb/fin regeneration-competent species.

In both the axolotl limb and zebrafish fin, amputation initiates a sequential activation of pro- and anti-inflammatory programs^15,49,50,82^. Similarly, our snRNA-seq analysis of the myeloid cluster indicates that following fin amputation, both pro- and anti-inflammatory signals contribute to the wound healing response. As regeneration progresses, anti-inflammatory signals become predominant (Fig. 8). Altogether, our findings provide further support for this wound healing dynamics as an evolutionarily shared feature of appendage regeneration^82^.

We leveraged our *Polypterus* and axolotl datasets to search for genes encoding candidate signaling factors expressed in the wound epidermis and blastema during fin and limb regeneration (Fig. 9). Our comparisons identified two genes highly expressed in the fin and limb wound epidermis, *tes* and *tmem154*. While little is known regarding cellular roles of *tmem154*, *tes* is a well-studied gene that functions as a tumor suppressor in the mouse^83^, and is positively associated with cell spreading^56^. Chicken cells overexpressing TES show increased cell spreading and decreased cell motility^84^. Given that cell spreading increases the contact surface of the cell with the extracellular matrix and its neighboring cells, upregulation of *tes* may help establish the integrity of the newly formed wound epidermis and its contact with the underlying extracellular matrix-rich blastema. In the connective tissue, we identified *tfpi2* and *tnfaip6* upregulation during fin and limb regeneration. TFPI-2, the product of *tfpi2*, is a serine proteinase inhibitor that helps regulate extracellular matrix digestion and remodeling^85^. In addition to being secreted, TFPI-2 translocates into the nucleus and interacts with AP-2a to negatively regulate the expression of MPP-2^86^. Furthermore, in cell culture studies, TFPI-2 upregulation promotes endothelial– mesenchymal transition (EndMT)^87^, a process that can contribute to myofibroblast generation. Extracellular matrix remodeling, regulation of metalloproteinase gene expression and EndMT are all processes fundamental for limb and fin regeneration. The versatile role played by TFPI-2 in these processes identifies it as a prime candidate for downstream functional studies. Single-cell RNA-seq analysis of zebrafish tail fin regeneration identified *tnfaip6* as one of the top differentially expressed genes in mesenchymal cells at 1 dpa^68^. Similarly, we observed *tnfaip6* upregulation in connective tissue cells contributing to the fin blastema. Given its established role in regulating mesenchymal cell proliferation and differentiation in mammals^67^, along with its upregulation during zebrafish fin regeneration and our findings of *tnfaip6* expression in both the *Polypterus* fin and axolotl limb blastema, *tnfaip6* may perform a conserved, evolutionarily shared function in vertebrate limb and fin regeneration.

The identification of TREEs active during limb and fin regeneration will provide valuable insights into the evolutionary dynamics of gene regulatory networks governing regeneration. As a first step towards this goal, our chromatin accessibility profiling of uninjured and regenerating fins at 3 dpa uncovered hundreds of differentially accessible peaks. Notably, many of the most significant peaks were located near genes that were upregulated during *Polypterus* fin regeneration such as *lima1*, *trim71*, and *tes* (Fig. 10). Like *tes*, the mammalian LIMA1 has been proposed to act as a mechanosensitive regulator of cell adhesion^88^, and both *lima1* and *tes* are upregulated in the wound epidermis of the *Polypterus* fin and axolotl limb. Trim71, along with Prdm16, Zbtb16, and Lin28, has been identified as part of a core set of four factors capable of successfully reprogramming mammalian non-limb fibroblasts into limb progenitor-like cells^72^. Peak 79, also among our top 100 most significant peaks, was found upstream of *bmp7a*, a gene implicated in limb regeneration^89^. Moreover, peak 18 was located upstream of *lepa*, the *Polypterus* ortholog to *lepb*, a gene sharply upregulated upon zebrafish fin ray amputation^60^. Expression of *lepb* during fin ray regeneration is regulated by a well-characterized TREE, termed *LEN*, located between *lebp* and mir129-1^60^. Interestingly, peak 18 was located over 1 mb upstream of the *Polypterus lepa* promoter, in a gene desert spanning nearly 3 mb between *mir129-1* and *snd1*. Hence, peak 18 is likely not an ortholog of LEN, but possibly an additional TREE, active in the context of fin amputation at the endoskeleton level. Finally, via differential TF footprinting, we identified an enrichment for predicted bound sites of AP-1 TFs during *Polypterus* fin regeneration, similar to recent reports in zebrafish^8,69^ and the axolotl^90^. Pending functional validation, the candidate TREEs identified here will represent a valuable resource to studies aimed at identifying shared and derived TREEs controlling limb and fin regeneration. A recent high-quality epigenetic profiling study of axolotl limb regeneration successfully identified and functionally validated an axolotl TREE^90^. However, since sampling consisted of fluorescence-activated cell sorting (FACS) of connective tissue cells, the axolotl dataset lacks the cellular complexity present in limbs and, consequently, is not expected to uncover TREEs driving gene expression in other limb tissues, including the wound epidermis. Ultimately, a comprehensive, high-quality epigenetic profiling of axolotl limb regeneration would provide the ideal dataset for comparative studies.

In conclusion, our study establishes *Polypterus* as a robust model that complements zebrafish by offering unique insights into the mechanisms underlying complete fin regeneration. Our findings highlight the power of comparative studies incorporating non-traditional research models to distinguish between ancestral and derived regenerative processes. Ultimately, a broad comparative approach that includes limb regeneration-incompetent species—such as post-metamorphic frogs, lizards, and mice—will be instrumental in identifying physiological, cellular, genetic, or epigenetic features that may have been modified or lost in amniotes.

## Materials and Methods

### Animal work

*Polypterus* (*P. senagalus*) and axolotls (*Ambystoma mexicanum*) specimens were maintained and used in accordance with an approved Louisiana State University (LSU) IACUC protocol IACUCAM-21-155. Juvenile fish were obtained from commercial vendors and maintained in individual tanks in a recirculating freshwater system at 27-28 °C and a day-night cycle as 12h light/12 h dark. Juvenile axolotls were obtained from the Ambystoma Genetic Stock Center at the University of Kentucky and maintained in individual tanks with 40% Holtfreter’s solution at 18-19 °C, and a day-night cycle as 12h light/12 h dark. All animals were anesthetized in 0.05% MS-222 (Sigma) before amputations. Pectoral fins of *Polypterus* fish ranging from 7 to 10 cm were bilaterally amputated across the endoskeleton using a sterile scalpel blade. For uninjured fin samples, a portion of the amputated fins encompassing the endoskeletal elements was sampled and processed according to downstream protocols. Regenerating fins were harvested at 1, 3 and 7 dpa. As we anticipate future comparisons of some of the sequence datasets described here with datasets generated from animals treated with specific protein inhibitors, regenerating fins were harvested from DMSO-treated animals (0.1% in fish system water) for bulk RNA-seq, spatial transcriptomics and snRNA-seq experiments. For axolotl samples, animals ranging from 8 to 12 cm had their forelimbs amputated at the level of the upper arm. Animals were euthanized according to our approved IACUC protocol (0.25% MS-222).

### Histology

Uninjured and regenerating tissues were flash-frozen using isopentane (2-methylbutane manufacturer) chilled with liquid nitrogen. Isopentane was poured into a metal canister and placed in liquid nitrogen to equilibrate. Tissues were embedded in Tissue-Tek O.C.T. compound (Sakura) in its appropriate mold. Using forceps, the molds containing O.C.T. compound were placed in the equilibrated isopentane. Samples were stored at -80°C for sectioning (10 μm thickness). Slides were stored at -80°C until subsequent use. Hematoxylin and Eosin staining was performed as previously described^3^.

### Hybridization Chain Reaction (HCR-FISH)

HCR was performed on 10 μm fresh frozen tissue sections following the manufacturer’s protocol (Molecular Instruments). Tissue sections were stored at −80°C until use. To begin, the sections were fixed in ice-cold 4% paraformaldehyde (PFA) (MP Biomedicals) in phosphate-buffered saline (PBS) for 15 minutes at room temperature. After fixation, the slides were washed three times with 1X PBS for 5 minutes each. Pre-hybridization was then performed at 37°C for 10 minutes. Probe hybridization was carried out overnight at 37°C. Following the hybridization, the slides were washed three times with probe wash buffer at 37°C for 15 minutes each, and then two washes were performed with 5X SSC-T buffer at room temperature for 5 minutes each. Pre-amplification was done by applying amplification buffer to the slides for 30 minutes at room temperature. Hairpins (H1 and H2) were denatured at 95°C for 90 seconds and allowed to cool to room temperature for 30 minutes in the dark. The hairpins were diluted 1:50 in amplification buffer and applied to each slide. The slides were incubated overnight at room temperature. After incubation, the slides were washed twice with 5X SSC-T buffer for 15 minutes each. ProLong Gold with DAPI (Invitrogen, #P36925) was used as mounting media and for nuclear counterstaining. Finally, the slides were stored at 4°C in the dark until imaging. Probes for this study were generated by Molecular Instruments and can be ordered using the following lot numbers: RTT023 (for *col6a3*) and RTT022 (for *col17a1b*).

### Bulk RNA-sequencing and analysis

For RNA-sequencing, total RNA was extracted from uninjured and 3 dpa pectoral fin tissue (3 biological replicates, 1 pair of fins per replicate) using TRIzol reagent (Invitrogen) according to the manufacturer’s protocol, followed by DNase I treatment and column clean-up (Qiagen RNeasy kit) to remove any residual DNA. mRNA enriched libraries were generated and sequenced at Novogene (https://www.novogene.com/us-en). Briefly, 1 μg of RNA per sample was used to generate sequencing libraries using the NEBNext Ultra RNA Library Prep Kit for Illumina, following the manufacturer’s recommendation. The sequencing was performed on an Illumina Novaseq 6000 platform to produce 150bp paired-end reads. Sequence reads were mapped to *Polypterus senegalus* Genome Assembly ASM1683550V1 using STAR V2.7.10b^91^. Read count extraction was performed using featureCounts v2.0.6^92^. Read count normalization and differential gene expression analysis were performed using DEseq2 R package^93^. For subsequent sequence analysis, the following list of genes were manually annotated according to sequence similarity and/or synteny to other vertebrate genomes: *alas2* (LOC120542955), *esam* (LOC120535937), *fcgbp-like* (LOC120541722), *mb1* (LOC120537046), *hif4a* (LOC120539324), *spp1* (LOC120527442), *tnfsf15* (LOC120535120), *tgfb1* (LOC120538772), *hs3st1l1* (LOC120540151), *clec7a-like* (LOC120535657), *timp1* (LOC120542910), *fn1* (LOC120530689), *pthlh-like* (LOC120538629), *fgf10* (LOC120527274), *krt19-like* (LOC120518116), *lima1* (LOC120524906), *tnc* (LOC120542391), *apoe-like* (LOC120541711).

### Spatial Transcriptomics

Cryosections (10 μm thickness) from regenerating pectoral fins (*Polypterus*) and regenerating forelimbs (axolotls) at 3 dpa were placed onto a 10X Genomics Visium Spatial Gene Expression Slide. Visium slide samples were processed using the 10X Genomics Spatial Gene Expression kit (#1000188) according to the instructions in the Visium Spatial Gene Expression Use Guide. Tissue permeabilization was performed for 18 minutes as previously determined using a 10X Genomics Visium Tissue Optimization Slide (#1000193). Brightfield histology images were obtained on a Leica DM6B Microscope at 20X magnification and stitched using the LAS X Office software. Generated libraries were sequenced at Novogene on an Illumina NovaSeq 6000 platform (paired-end, 150bp), with the sequencing depth of 50,000 reads per tissue covered spot. Raw sequence data (FASTQ) and histology images were processed using Space Ranger 3.0.1. Data visualization and generation of specific gene expression images were performed using Loupe Browser 8.0.0.

### Nuclei preparation and single-nucleus RNA-sequencing

Uninjured and regenerating pectoral fins at 1, 3 and 7 dpa were dissected using sterile scalpel blades and spring scissors. For each time point, a pair of pectoral fins from two individuals was collected and pooled as one sample after nuclei isolation. A total of two biological replicates (each comprising two polled animal samples) were prepared for each time point. Pectoral fin tissue from each animal was minced first with a spring scissor and then with a razor blade in a petri dish on ice. The minced tissue was resuspended in 1 mL of 0.3X lysis buffer (Active Motif ATAC-seq lysis buffer diluted 1:3 with a lysis dilution buffer: 10 mM Tris-HCl, pH: 7.5, 50 mM NaCl, 20 mM Mg2Cl, 10 % BSA) and then homogenized in a pre-chilled Dounce homogenizer (12 strokes). The homogenate was filtered using a 40 μm cell strainer and immediately diluted with additional 4 mL of lysis dilution buffer and then centrifuged at 400 xg at 4 °C for 5 minutes in a swinging bucket rotor. The pellet was quickly washed with 1 mL of wash buffer (1X PBS/0.5% BSA), containing 40U/μl of Protector RNAse inhibitor (Millipore-Sigma #3335399001), without resuspension and then resuspended in 0.5 mL of wash buffer. At this point, samples from the two individuals were combined. Nuclei were counted using a hemocytometer and then a volume corresponding to 10^6^ nuclei was transferred to an Eppendorf LoBind microcentrifuge tube and centrifuged at 400 xg at 4 °C for 5 minutes. Nuclei were fixed and frozen using the Evercode Nuclei Fixation Kit v3 from Parse Biosciences following the manufacturer’s protocol. Fixed and frozen nuclei were then processed to generate barcoded libraries aiming for 7,000 nuclei per sample using the Evercode WT Mini v3 Kit as per manufacturer’s instructions. Resulting libraries were sequenced targeting 20,000 reads per nuclei in an Illumina Novaseq Xplus platform (paired-end, 150bp).

### Single-nucleus RNA-seq analysis

SnRNA-seq FASTQ files were processed using the Trailmaker^TM^ piping module (https://app.trailmaker.parsebiosciences.com/; pipeline v1.4.0, Parse Biosciences, 2024). In this module sequences were processed for demultiplexing, barcode correction, read alignment and transcript quantification. Quantified transcripts were used to generate a cell-by-gene count matrix that was automatically redirected to the Insight module of Trailmaker for downstream analysis. During the Insight module processing, background, doublets and poor-quality nuclei (high content of mitochondrial gene detection and low level of detected transcripts) were excluded using the Trailmaker default data processing. For our dataset the minimum number of transcripts per cell varied from 390 to 928, and the median number of transcripts per cell after all filters ranged from 1909 to 5199. The maximum cutoff value for maximal percentage of mitochondrial genes was 1.21. Data normalization, principal-component analysis (PCA) and data integration using Harmony were performed on data from high-quality cells. Clusters were identified using the Leiden method, and a Uniform Manifold Approximation and Projection (UMAP) embedding was calculated to visualize the results. Cluster-specific marker genes were identified by comparing cells of each cluster to all other cells using the presto package implementation of the Wilcoxon rank-sum test. Seurat v5 R package^94^ was used to produce heatmaps and gene expression UMAP image panels.

### Bulk ATAC-seq library preparation

Uninjured and 3 dpa fin tissues were collected and subsequently used for nuclei isolation and tagmentation following the ATAC-seq kit protocol (Active Motif, #105320) without modifications. For each triplicate, tissue from both left and right pectoral fins were collected and pooled to comprise one biological replica. Each library was prepared using 50,000 nuclei and amplified within ten PCR cycles. After PCR amplification, the libraries were purified and sent for sequencing. The sequencing was carried out using the NovaSeq XPlus platform, generating 150bp paired-end reads (150 million reads/sample).

### ATAC-seq data analysis

The three biological replicates of each condition were processed as follows. Raw reads were trimmed using NGmerge (version 0.3)^95^. The trimmed reads were aligned to the *Polypterus senegalus* reference genome from NCBI (ASM1683550v1) using Bowtie2 (version 2.2.5)^96^. Alignments were output as a BAM file, indexed using Samtools (version 1.18)^97^. Fragment size distribution was assessed using the ATACseqQC package (version 1.24.0)^98^ in RStudio (version 4.3.1). Trimmed reads were name-sorted using Samtools (version 1.18)^97^. Genrich (version 0.6.1) was used to call peaks using the parameters -j -y -r -v -m -e -f, to remove duplicate reads, multimapped reads, and reads mapped to mitochondrial chromosome. Downstream analysis of the ATAC-seq data was performed using DiffBind (version 3.10.1) to identify differential chromatin accessibility (peaks) between uninjured and regenerating fin tissues^99^. Peak sets generated by Genrich were associated with metadata for each sample and imported into DiffBind within RStudio (version 4.3.1). Read counts were generated using summits = 75 and normalized using DESeq2 (version 1.40.2)^100^. Peaks that exhibited differential accessibility between the two conditions were selected based on a significance threshold of FDR < 0.05. Peaks were visualized using Integrative Genomics Viewer (IGV) (version 2.14.1). AnnotatePeaks.pl from HOMER (version 4.11) was used to identify and annotate the genes located nearest to peaks. Motif footprint enrichment was performed using TOBIAS^74^.

## Acknowledgments

We thank Patricia Schneider for helpful feedback and comments. We also thank Jamily Lima for creating drawings that contributed to the final illustrations. This work was funded by start-up funds from Louisiana State University (I.S.), and an NSF-Integrative Organismal Systems (IOS) Enabling Discovery through GEnomics (EDGE) grant (2421117, I.S.).

---

**Supplementary Fig. 1.**
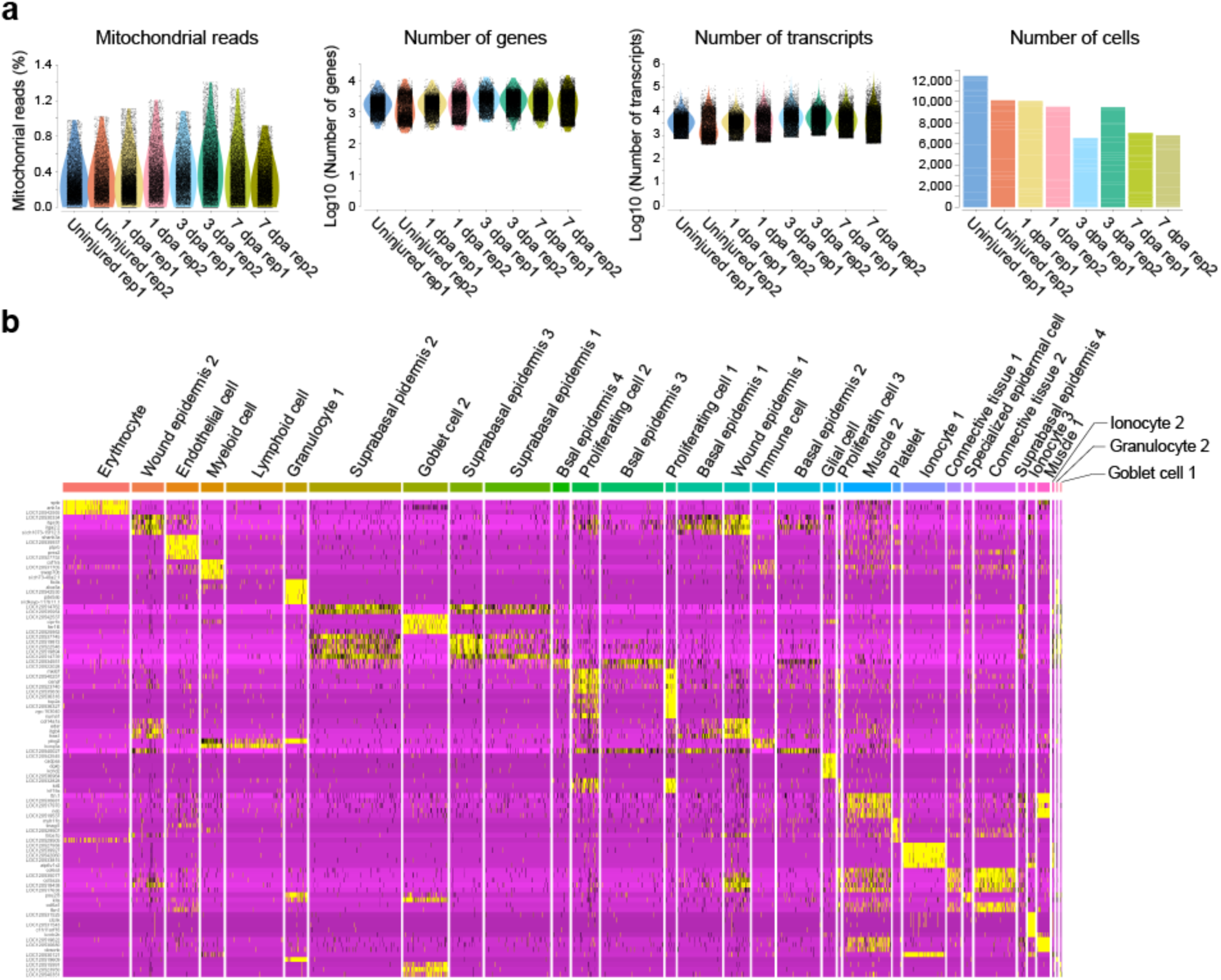
*Polypterus* fin snRNA-Seq quality control metrics and marker gene expression across different cell clusters. **a** Percentage of mitochondrial reads, number of genes, transcripts and cells (nuclei) across samples; samples are color-coded as indicated above the figure. **B** Heatmap depicting marker gene expression across different cell clusters.

**Supplementary Fig. 2.**
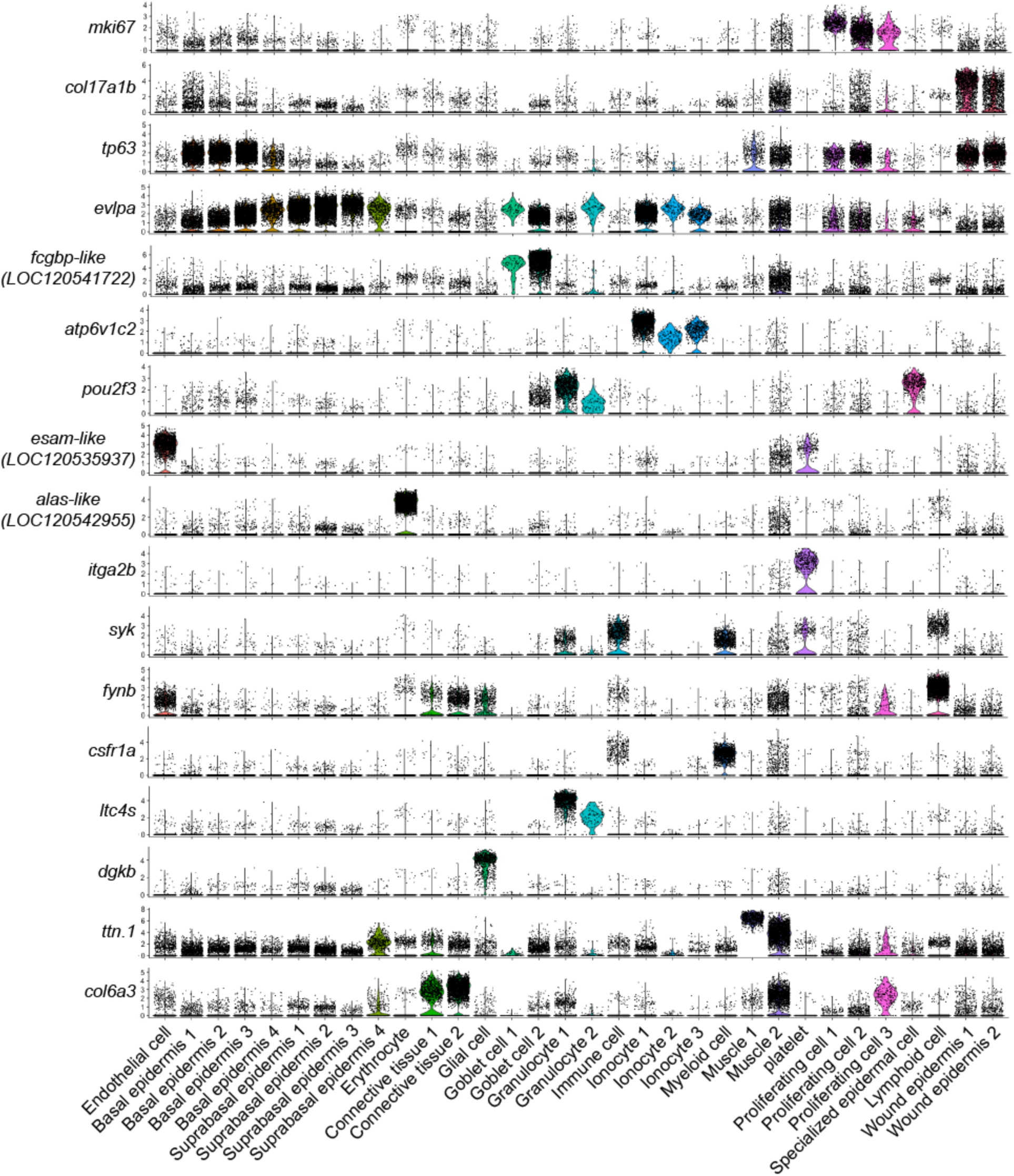
Marker gene expression of major cell populations across all stages. Violin plots of select markers of major cell populations present in all stages (uninjured, 1 dpa, 3 dpa, and 7 dpa).

**Supplementary Fig. 3.**
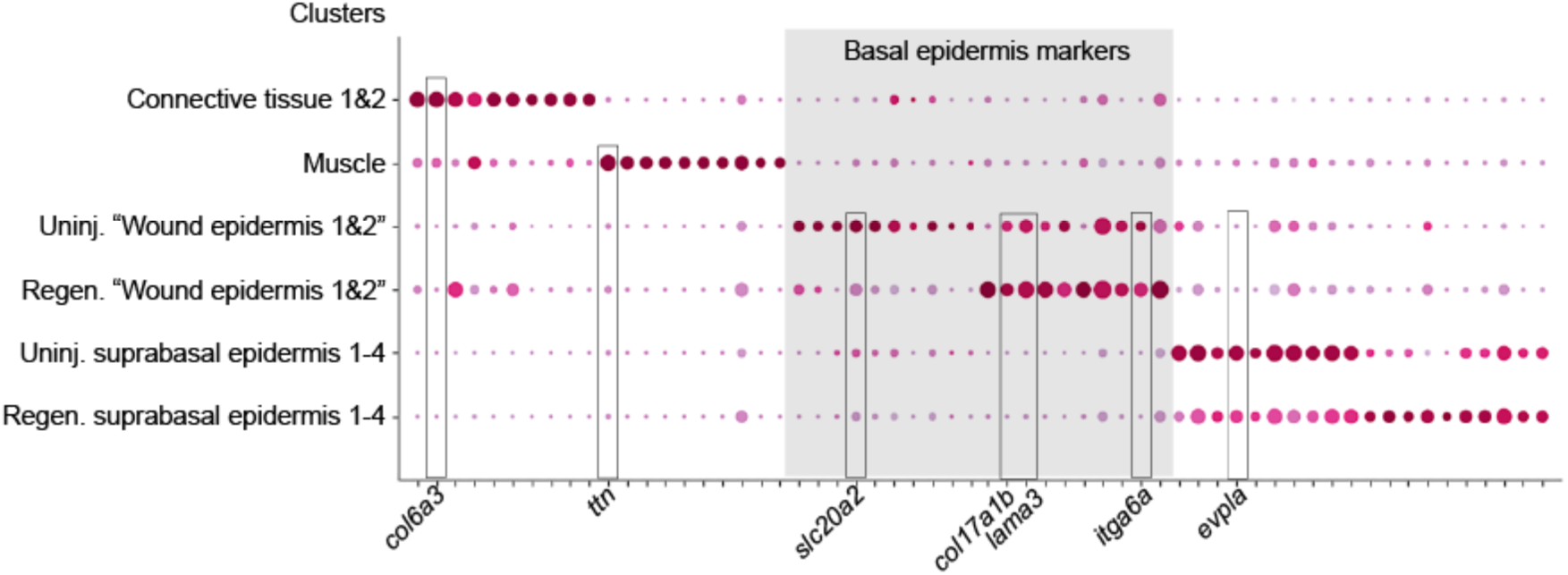
Cells annotated as wound epidermis at homeostasis express basal epidermis cell markers. Dot plot shows that “wound epidermis” cells in the uninjured fin express shared basal epidermis markers with wound epidermis clusters 1 and 2 of regeneration stages (1 dpa, 3 dpa, and 7 dpa).

**Supplementary Fig. 4.**
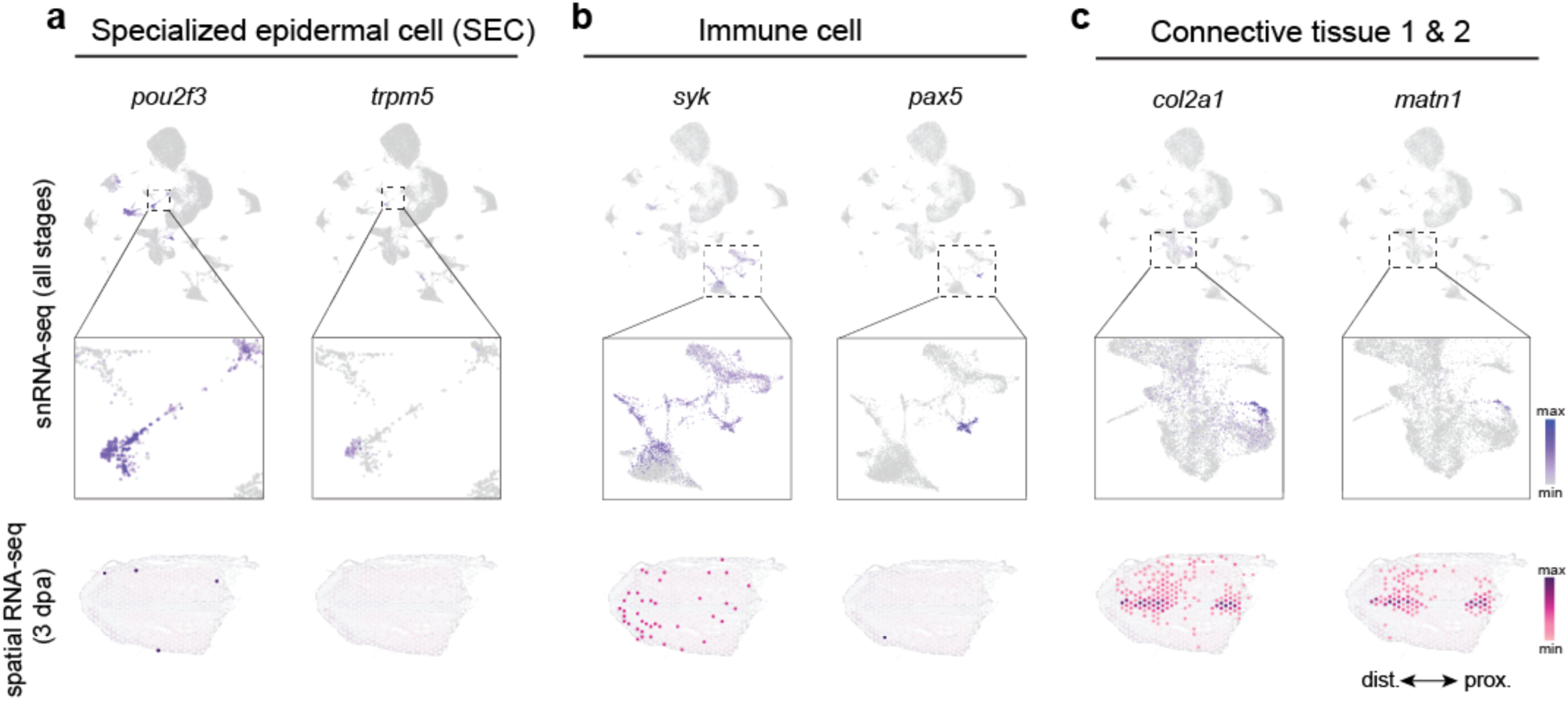
Spatial and snRNA-seq helps refine annotation of clusters and select cell type. **a** Expression of *pou2f3* and *trpm5* in the specialized epithelial cell cluster; *pou2f3* was detected in a few spatial RNA-seq spots in the epidermis at 3 dpa, whereas *trpm5* was not detected. **b** Expression of *syk* and *pax5* in the immune cell cluster; *syk* was detected in scattered spots, mostly distal in spatial RNA-seq in the wound epidermis and blastema region at 3 dpa, whereas *pax5* was detected in a single spot in the presumptive wound epidermis. **c** Expression of *col2a1* and *matn1* in a subset of cells in the connective tissue clusters 1 and 2; *col2a1* and *matn1* were detected in spots along the fin skeleton at 3 dpa.

**Supplementary Fig. 5.**
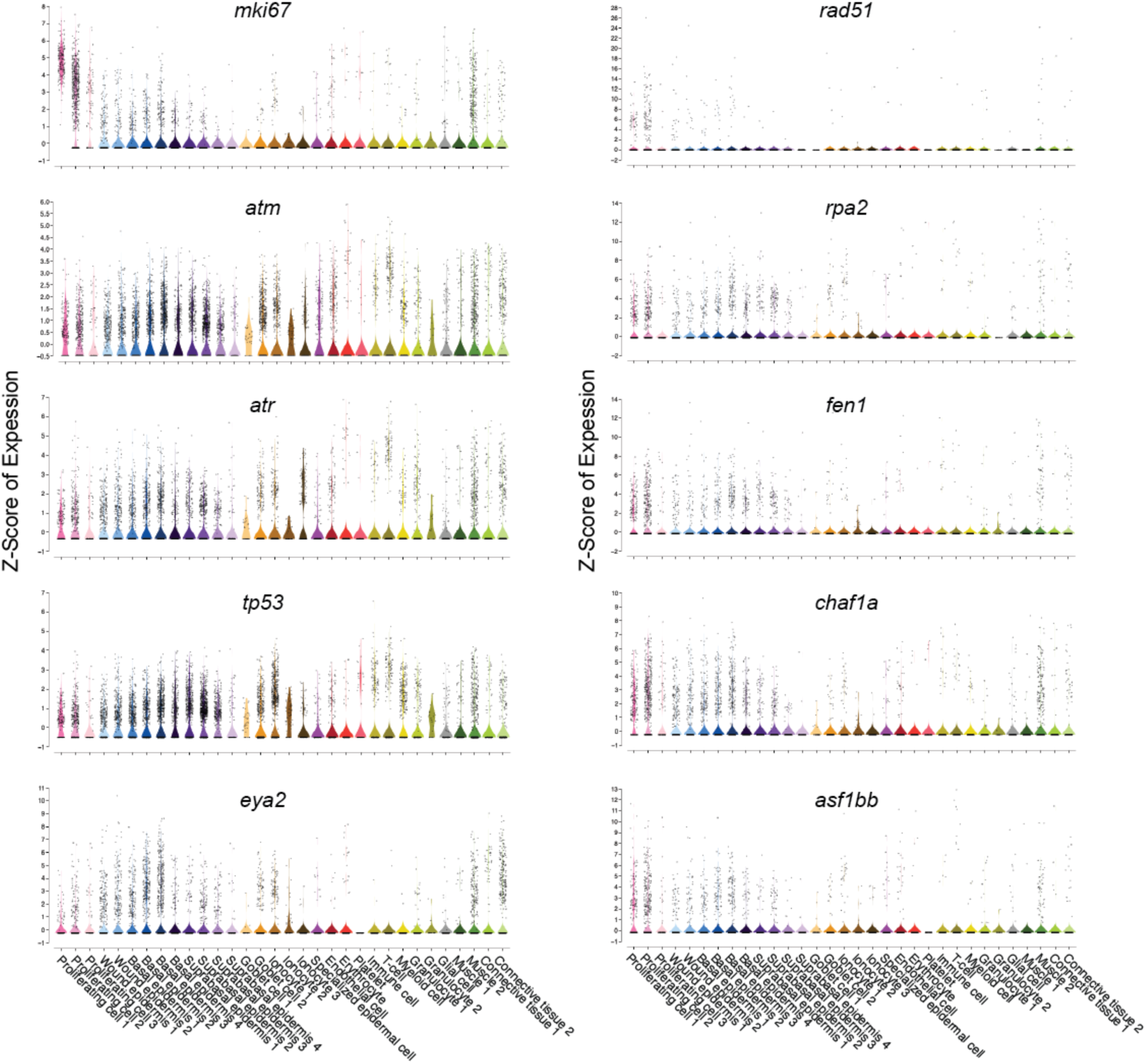
Spatial and snRNA-seq helps refine annotation of clusters and select cell type. Violin plots showing expression markers of cell proliferation, DNA damage sensing and DNA damage repair in snRNA-seq clusters.

**Supplementary Fig. 6.**
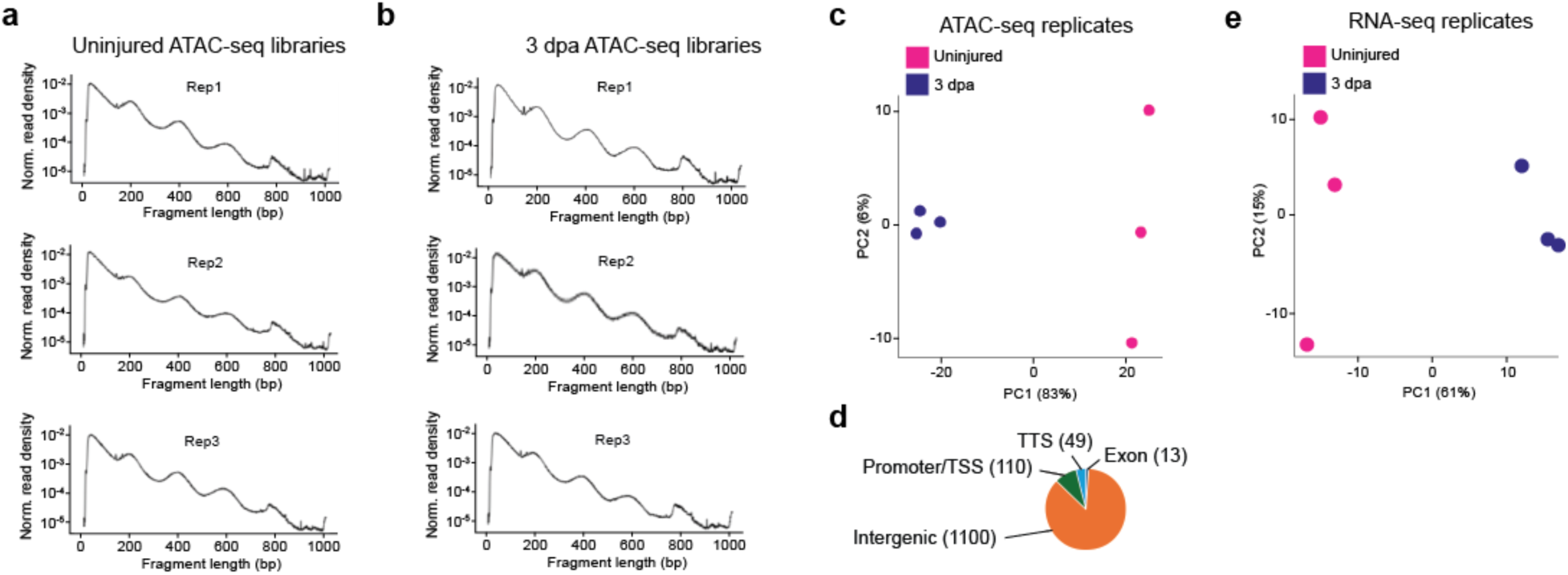
Quality control of bulk ATAC-seq and bulk RNA-seq libraries. Fragment distribution of ATAC-seq aligned BAM files from uninjured (**a**) and 3 dpa stage (**b**) fin libraries in logarithmic scale. **c** Principal component analysis showing separation between ATAC-seq replicates. **d** Annotation of ATAC-seq peaks; number of peaks per category shown in parenthesis. **e** Principal component analysis showing separation between RNA-seq replicates.

**Supplementary Fig. 7.**
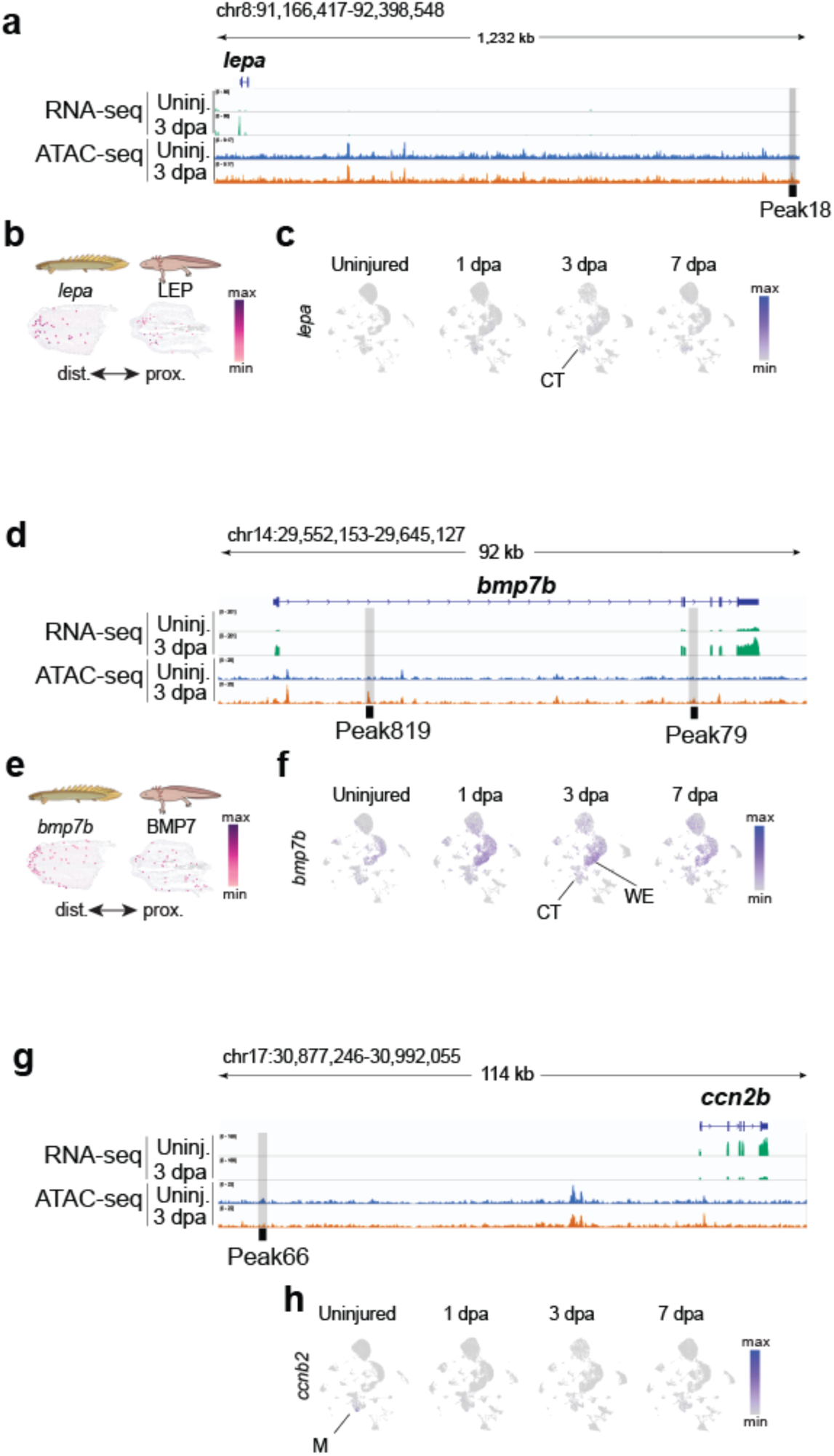
Additional examples of *Polypterus* candidate TREEs. Genomic location of peaks 18 upstream of *lepa* (**a**), peaks 79 and 819 upstream of *bmp7b* (**d**), and peak 66 upstream of *ccn2b* (**g**), including bulk RNA-seq and ATAC-seq tracks. Spatial expression of *Polypterus lep* (**b**), *bmp7b* (**e**), and their respective axolotl orthologs at 3 dpa. SnRNA-seq UMAP showing expression of *lepa* (**c**), *bmp7b* (**f**), and *ccn2b* (**i**). M, muscle; CT, connective tissue; WE, wound epidermis.

